# CD33^KO^-CD33-mesothelin loop CAR design avoids fratricide and improves efficacy of iNK cells against acute myeloid leukemia

**DOI:** 10.1101/2025.01.23.634500

**Authors:** Yao Wang, Xiujuan Zheng, Zhiqian Wang, Ziyun Xiao, Yunqing Lin, Fan Zhang, Yanhong Liu, Pengcheng Liu, Qitong Weng, Leqiang Zhang, Chengxiang Xia, Dehao Huang, Lijuan Liu, Yanping Zhu, Qi Zhang, Hanmeng Qi, Yi Chen, Yiyuan Shen, Chenyuan Zhang, Jiacheng Xu, Yaoqin Zhao, Jiaxin Wu, Tongjie Wang, Mengyun Zhang, Minming Li, Wenbin Qian, Aibin Liang, Xin Du, Wenyu Yang, Qi Chen, Xiaofan Zhu, Fangxiao Hu, Jinyong Wang

**Affiliations:** Guangzhou Institutes of Biomedicine and Health, Chinese Academy of Sciences, Guangzhou 510530, China; State Key Laboratory of Organ Regeneration and Reconstruction, Institute of Zoology, Chinese Academy of Sciences, Beijing 100101, China; Beijing Institute for Stem Cell and Regenerative Medicine, Beijing 100101, China; University of Chinese Academy of Sciences, Beijing 100049, China; Department of Hematology, Guangdong Provincial People’s Hospital, Guangdong Academy of Medical Sciences, Guangzhou 510080, China; Department of Hematology, the Second Affiliated Hospital, Zhejiang University School of Medicine, Hangzhou 310009, China; Department of Hematology, Tongji Hospital of Tongji University, Shanghai 200065, China; State Key Laboratory of Experimental Hematology, National Clinical Research Center for Blood Diseases, Haihe Laboratory of Cell Ecosystem, Institute of Hematology & Blood Diseases Hospital, Chinese Academy of Medical Science & Peking Union Medical College, Tianjin 300020, China; Center for cell lineage and development, CAS Key Laboratory of Regenerative Biology, Guangdong Provincial Key Laboratory of Stem Cell and Regenerative Medicine, GIBH-HKU Guangdong-Hong Kong Stem Cell and Regenerative Medicine Research Centre, GIBH-CUHK Joint Research Laboratory on Stem Cell and Regenerative Medicine, Guangzhou Institutes of Biomedicine and Health, Chinese Academy of Sciences, Guangzhou, 510530, China

## Abstract

Acute myeloid leukemia (AML) patients are often older, which brings challenges of endurance and persistent efficacy of autologous CAR-T cell therapies. Allogenic CAR-NK cell therapies may offer reduced toxicities and enhanced anti-leukemic potential against AML. In this study, we designed a novel CD33-mesothelin loop CAR (Loop CAR) and evaluated its anti-tumor efficacy in human umbilical cord blood-derived NK (UCB-NK) cells and human pluripotent stem cell-derived NK (hPSC-iNK) cells. The Loop CAR exhibited superior cytotoxicity against dual-antigen-positive tumor cell lines and primary AML cells. To further avoid fratricide caused by endogenous CD33 expression in NK cells, we established a hPSC-derived cell line via knockout of CD33 gene (CD33^KO^) and engineered Loop CAR. Rather than enforced expression of exogenous CD16, we generated abundant mature CD33^KO^-Loop CAR-iNK cells highly expressing endogenous CD16 via an organoid induction approach. This innovative strategy effectively mitigated NK cell fratricide and significantly enhanced CD33 and mesothelin-mediated specific cytotoxicity. Moreover, the CD33^KO^-Loop CAR-iNK cells demonstrated superior tumor-killing activity in AML xenograft mice and significantly prolonged survival. hPSC-derived CD33^KO^-Loop CAR-iNK cells possess unique advantages and translational potential for treating AML.

## INTRODUCTION

Acute myeloid leukemia (AML) is an aggressive hematologic malignancy that demands urgent therapy (*1*). As new therapeutic targets are being identified, significant efforts have highlighted mesothelin (MSLN) as a promising target in AML. The MSLN is expressed in 36% of pediatric AML cases and 14% of adult AML cases, with 76% of MSLN-positive (MSLN^+^) patients retaining MSLN expression at relapse. Furthermore, 4% of MSLN-negative (MSLN^-^) AML patients acquire MSLN expression upon relapse (*2–4*). Preclinical data suggest that MSLN chimeric antigen receptor (CAR)-T cells effectively eliminate MSLN^+^ AML cells in xenograft models (*5*). Therefore, MSLN is a promising target for AML immunotherapy.

Natural killer (NK) cells have demonstrated considerable potential in enhancing outcomes for AML treatment, particularly in the context of hematopoietic stem cell transplantation (*6*). The adoptive transfer of NK cells for AML therapy has been widely investigated in clinical settings (*7*). CAR-NK cells, which combine the target killing ability of CAR technology with the innate nonspecific cytotoxicity of NK cells, have shown promising results for tumor treatment (*8, 9*). CD33 CAR-NK cells, in particular, have been tested in relapsed/refractory AML therapies, with clinical data demonstrating their efficacy and safety (*10*). Additionally, pluripotent stem cell (PSC)-derived NK or CAR-NK cells have shown therapeutic potential in both preclinical and clinical settings for AML (*11, 12*). PSC-derived NK (iNK) cells provide a homogeneous alternative to NK cells isolated from human tissues (*11, 13, 14*).

Despite these advancements, challenges remain in CAR immune cell therapy, such as tumor antigen escape, on-target off-tumor toxicity, and unintended fratricide due to the shared expression of target antigens between CAR immune cells and tumor cells (*15–17*). To overcome these issues, dual-targeting CARs have been designed to simultaneously target two antigens expressed on tumor cells (*18, 19*). Inactivation of the target antigen in the CAR-engineered immune cells, or in healthy tissue cells, can mitigate fratricide and spare normal tissues. For example, the disruption of CD7 in CD7 CAR-T cells prevents fratricide and allows for the expansion of these cells (*20*). Similarly, genetic inactivation of CD33 in hematopoietic stem cells enables CD33-deficient hematopoiesis to avoid the killing of CD33 CAR-T cells (*21*). Since CD33 is also expressed on NK cells, CD33 CAR-NK cells face similar fratricidal challenges, which can hinder their expansion (*17, 22*). Human umbilical cord blood NK (UCB-NK) cells, which exhibit a 1.4%-25.5% CD33 expression range in their resting state, show reduced cytotoxicity against K562 cells (*23*). Moreover, NK cells upregulate CD33 expression levels to nearly 50% during *in vitro* expansion (*17*).

In this study, we designed a bivalent loop CAR targeting both CD33 and MSLN antigens (CD33-MSLN loop CAR, denoted as Loop CAR). We engineered this Loop CAR into UCB-NK cells and hPSCs to generate Loop CAR-NK and Loop CAR-iNK cells. Both Loop CAR-NK and Loop CAR-iNK cells exhibited superior cytotoxicity against tumor cells expressing CD33 and MSLN antigens, including AML cell lines and patient-derived tumor cells. Transcriptomic analysis confirmed that the Loop CAR enhanced NK cell cytotoxicity. Additionally, we generated a CD33 gene knockout (CD33^KO^) hPSC line expressing the Loop CAR (CD33^KO^-Loop CAR-hPSCs) and successfully produced CD33^KO^-Loop CAR-iNK cells. These CD33^KO^-Loop CAR-iNK cells exhibited enhanced expansion and anti-tumor efficacy *in vitro* and effectively suppressed tumor progression in xenograft AML models. Our findings highlight the clinical potential of Loop CAR-NK and Loop CAR-iNK cells for the treatment of CD33^+^MSLN^+^ AML.

## RESULTS

### CD33-MSLN loop CAR-NK cells exhibit superior cytotoxicity against CD33- and MSLN-positive tumor cells

To enhance the efficacy of NK cells targeting mesothelin (MSLN)-expressing acute myeloid leukemia (AML), we employed a bivalent loop CAR strategy, designing a dual-target CAR targeting both CD33 and MSLN. Specifically, the variable light (V_L_) and heavy (V_H_) chains of anti-CD33 single-chain variable fragment (CD33 scFv) and anti-MSLN scFv (MSLN scFv), the (G_4_S)_3_ linker (Linker 3), and the Whitlow linker (*24*) were sequentially fused to create a bivalent scFv construct (CD33 V_L_-linker 3-MSLN V_H_-whitlow linker-MSLN V_L_-linker 3-CD33 V_H_, CD33-MSLN loop). The CD33-MSLN loop scFv was then linked to the CD8α hinge, the CD8 transmembrane domain (TMD), and the CD3 ζ signaling domain (SD) to construct the final CD33-MSLN loop CAR (denoted as Loop CAR). We applied the MSLN CAR and CD33 CAR as controls (**Fig. 1A**). To assess the potential of Loop CAR in enhancing tumor-killing activity in NK cells, we first incorporated the Loop CAR element into the SFG vector, followed by retroviral packaging as previously reported (*25*). After T-cell depletion, umbilical cord blood NK (UCB-NK) cells were activated using K562-mbIL-21 feeder cells for 6 days, followed by transduction with Loop CAR retroviral particles (MOI = 5). The transduced cells were then expanded for 4 days in the presence of K562-mbIL-21 feeder cells (**Fig. 1B** and **fig. S1A**). The proportion of Loop CAR^+^ NK cells (CD56^+^CD33 CAR^+^MSLN CAR^+^) reached 43.4% two days after transduction, while control groups, including EGFP-NK, CD33 CAR-NK, and MSLN CAR-NK, achieved transduction rates of 48.2%, 55.6%, and 52.1%, respectively (**Fig. 1C**). Loop CAR-NK cells showed comparable expression of typical NK cell activating and inhibitory receptors, as well as cytotoxic granules, to the control groups (**fig. S1B**).

**Fig. 1.**
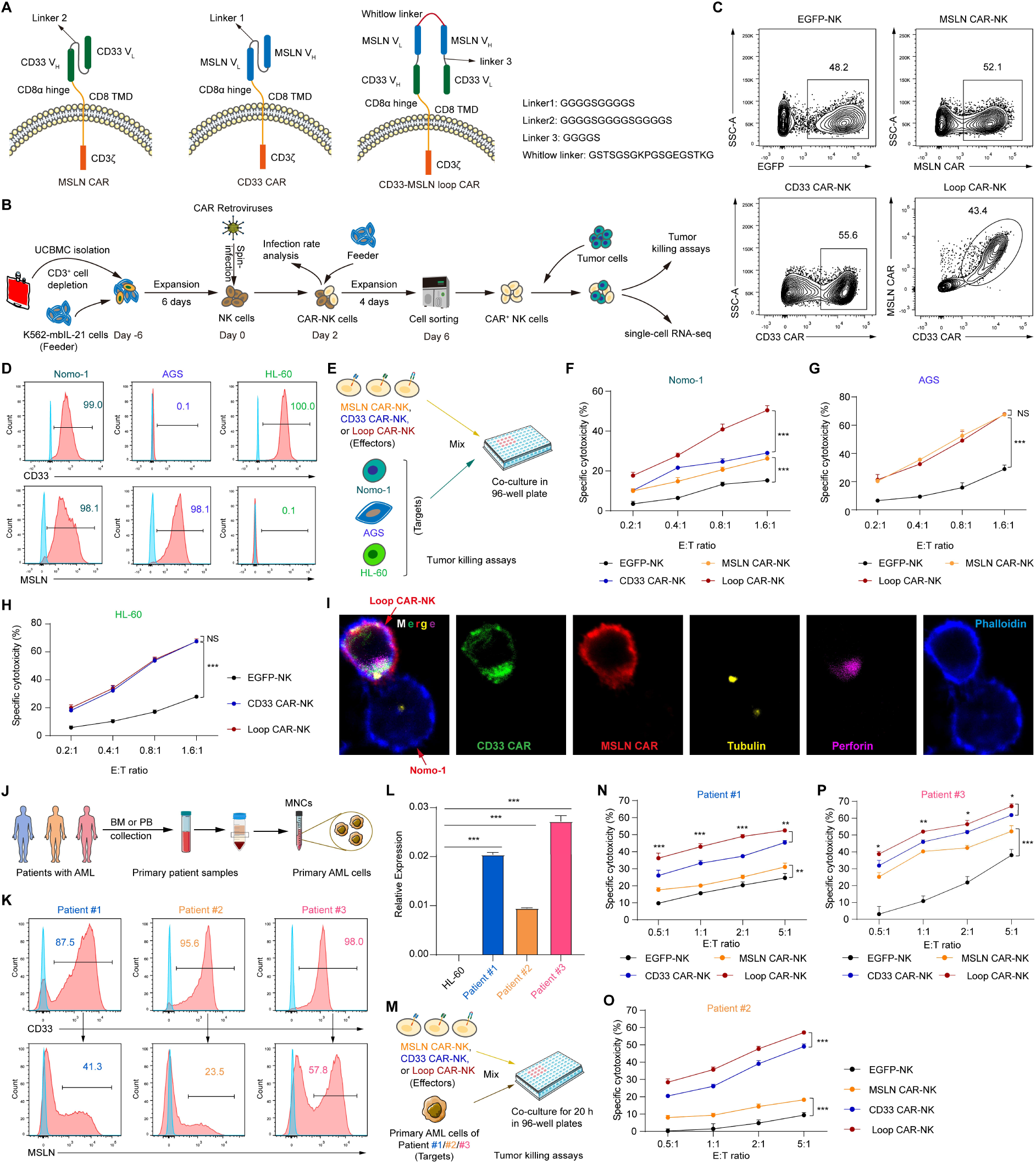
Development and functional evaluation of Loop CAR-NK cells. (**A**) Schematic diagram of the design of Loop CAR. (**B**) Schematic diagram showing the generation of Loop CAR-NK cells procedures and evaluation their cytotoxicity. (**C**) Flow cytometric analysis of the transduction efficiencies of each group of CAR-NK cells. EGFP virus-infected NK cells were used as control. (**D**) Flow cytometry histograms showing the expression levels of CD33 and MSLN in target cell lines, including CD33^+^MSLN^+^ Nomo-1 AML cells, CD33^-^MSLN^+^ AGS gastric adenocarcinoma cells, and CD33^+^MSLN^-^ HL-60 AML cells. (**E**) Tumor killing assay design. (**F-H**) Statistics of CAR-NK cells cytotoxicity against Nomo-1 tumor cells (F), AGS cells (G), and HL-60 cells (**H**) at indicated E:T ratios. Nomo-1 cells and HL-60 cells were incubated with effectors for 4 hours (n = 4 per group). AGS cells were incubated with effectors for 6 hours (n = 4 per group). (**I**) Confocal microscopy representative images of immunological synapse formed between loop CAR-NK cells and Nomo-1 cells. The conjugates were stained with antibodies against perforin (Violet), phalloidin-F-actin (Blue), CD33 CAR (Green) and MSLN CAR (Red). (**J**) Schematic diagram showing the collection of peripheral blood or bone marrow cells from patients with AML. (**K**) Flow cytometric analysis of the expression of CD33 and MSLN in primary AML cells. (**L**) Real-time quantitative PCR analysis of MSLN transcript expression levels in primary AML cells. The *GAPDH* gene was used as an internal control. Relative gene expression levels were calculated as 2^−ΔCt^. (**M**) Schematic diagram showing primary AML cells killed by CAR-NK cells *in vitro*. Each group of NK cells was co-cultured with primary AML cells for 20 hours at indicated E:T ratios (*n* = 4 per group). (**N-P**) Specific cytotoxicity of CAR-NK cells derived from UCB-NK against primary AML cells from three patients (Patient #1: N, Patient #2: O, Patient #3: P). Statistical analysis: one-way ANOVA. NS: not significant; \**P* < 0.05, \*\**P* < 0.01, \*\*\**P* < 0.001.

We next evaluated the cytotoxicity of Loop CAR-NK cells by sorting CAR^+^ NK cells and coculturing them with Nomo-1 (CD33^+^MSLN^+^), AGS (CD33^-^MSLN^+^), or HL-60 (CD33^+^MSLN^-^) tumor cells (**Fig. 1**, **D** and **E**, **fig. S1C**). Loop CAR-NK cells exhibited significantly superior cytotoxicity against Nomo-1 cells compared to EGFP-NK, CD33 CAR-NK, and MSLN CAR-NK cells across all effector-to-target (E:T) ratios (E:T = 0.2:1, 0.4:1, 0.8:1, 1.6:1) (**Fig. 1F**). In addition, Loop CAR-NK cells also demonstrated potent killing activity against MSLN- or CD33-expressing tumor cells, such as AGS (CD33^-^MSLN^+^) and HL-60 (CD33^+^MSLN^-^), with cytotoxicity comparable to MSLN CAR-NK or CD33 CAR-NK cells (**Fig. 1**, **G** and **H**). Mechanistically, Loop CAR-NK cells exhibited the highest expression of CD107a, a marker associated with NK cell functional activity, when cocultured with Nomo-1 cells (**fig. S1**, **D-F**). Furthermore, Loop CAR molecules accumulated at the immunological synapse (IS) formed between Loop CAR-NK cells and Nomo-1 cells, confirming that the Loop CAR participates in IS formation in a CD33-MSLN-dependent manner (**Fig. 1I**). Notably, Loop CAR-NK cells also demonstrated superior cytotoxicity against primary AML cells expressing both CD33 and MSLN. Mononuclear cells (MNCs) were isolated from the bone marrow (BM) or peripheral blood (PB) of three AML patients (**Fig. 1J**). All three patients’ MNCs expressed CD33 (Patient #1: 87.5%, Patient #2: 95.6%, Patient #3: 98.0%) and MSLN (Patient #1: 41.3%, Patient #2: 23.5%, Patient #3: 57.8%) (**Fig. 1K**). The expression of MSLN was further confirmed by qPCR, which showed consistency with the protein expression levels (**Fig. 1L** and **table S1**). Loop CAR-NK cells (Effectors, E) were cocultured with these primary AML cells (Targets, T) for 20 hours at various E:T ratios (0:1, 0.5:1, 1:1, 2:1, and 5:1) (**Fig. 1M**). Loop CAR-NK cells exhibited significantly higher cytotoxicity against all patient-derived targets compared to CD33 CAR-NK and MSLN CAR-NK cells. In addition, both CD33 CAR-NK and MSLN CAR-NK cells killed more targets than EGFP-NK cells (**Fig. 1**, **N-P**). Collectively, these data demonstrate that Loop CAR-NK cells exhibit superior specific cytotoxicity against tumor cells expressing both CD33 and MSLN compared to CD33 CAR-NK or MSLN CAR-NK cells.

### Loop CAR-NK cells exhibit a more pronounced activation status after incubation with Nomo-1 cells

To investigate the superior cytotoxicity of Loop CAR-NK cells, we performed single-cell RNA sequencing (scRNA-seq) on sorted Loop CAR-NK cells after incubation with Nomo-1 cells for 72 hours (**Fig. 2A**). UMAP analysis revealed that the transcriptional profiles of Loop CAR-NK cells were similar to those of CD33 CAR-NK or MSLN CAR-NK cells. However, all CAR-NK cells exhibited distinct transcriptional profiles compared to EGFP-NK cells (**Fig. 2B**). We further examined the differentially expressed genes (DEGs) between Loop CAR-NK cells and CD33 CAR-NK or MSLN CAR-NK cells. Gene Ontology (GO) enrichment analysis indicated that genes upregulated in Loop CAR-NK cells relative to MSLN CAR-NK cells were primarily involved in cell division, positive regulation of the Fc receptor-mediated stimulatory signaling pathway, and positive regulation of the RIG-1 signaling pathway (**Fig. 2C** and **fig. S2**, **A** and **B**). Similarly, genes upregulated in Loop CAR-NK cells compared to CD33 CAR-NK cells were enriched in lymphoid cell differentiation, positive regulation of the RIG-1 signaling pathway, and formation of immunological synapses. (**Fig 2D** and **fig. S2**, **C** and **D**). Among the activation-related genes, several hallmarks of NK cell activation and cytotoxic function, including *CD244*, *SLAMF7*, and *ITGAL* (LFA-1), were upregulated (*26–28*). Additionally, genes involved in the RIG-I signaling pathway, such as *ANKRD17*, *PUM1*, and *PUM2* (*29, 30*), and those associated with immunological synapse (IS) formation, including *NCAM1*, *PTK2B*, and *DOCK8* (*31–34*), were also significantly elevated (**Fig. 2**, **E-G**). These findings suggest that Loop CAR-NK cells exhibit enhanced activation following incubation with Nomo-1 cells, which likely contributes to their superior cytotoxicity compared to CD33 CAR-NK or MSLN CAR-NK cells.

**Fig. 2.**
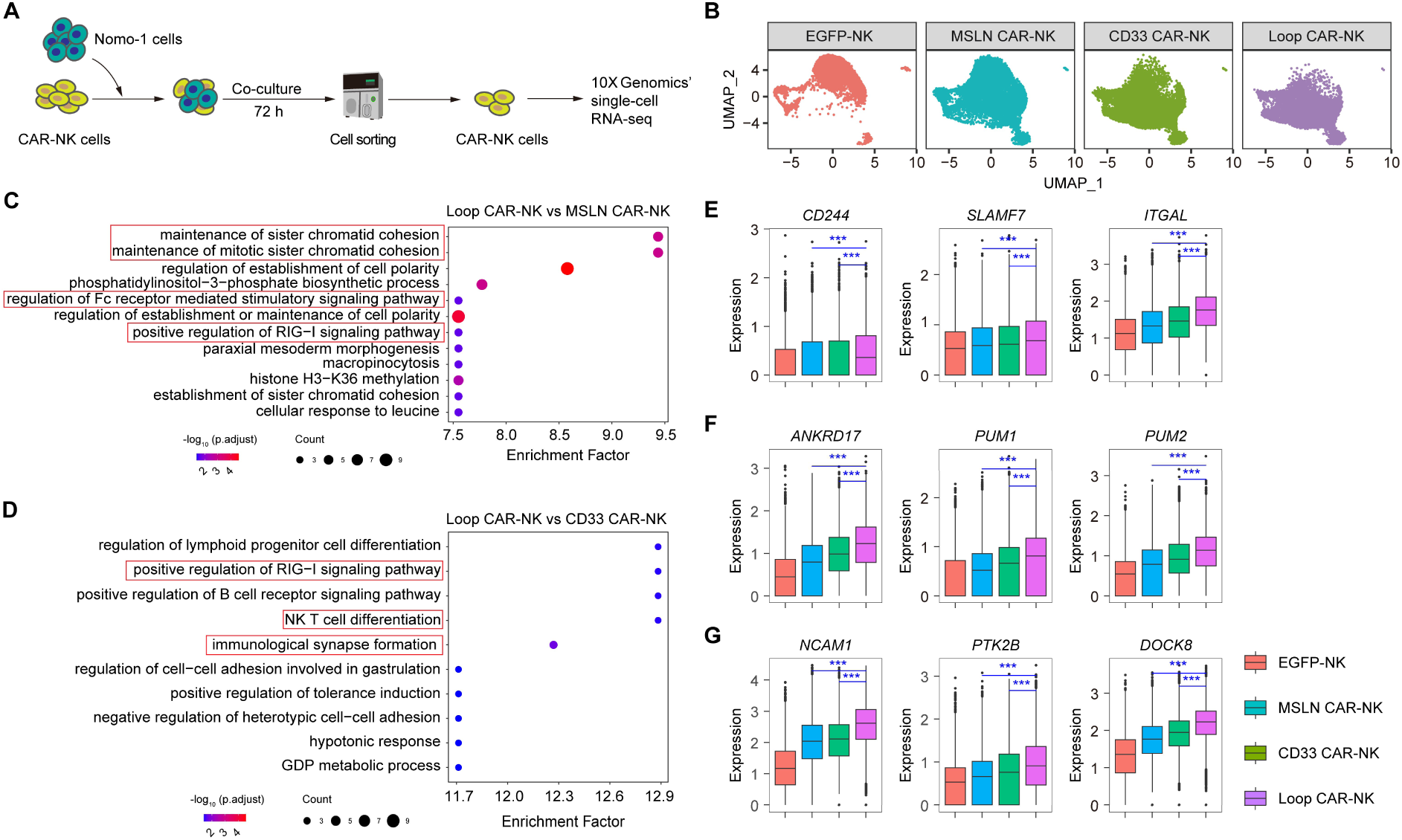
Molecular characterization of Loop CAR-NK cells. (**A**) Schematic diagram of sorting CAR-NK cells for 10× Genomics’ single-cell RNA-seq assay. CAR-NK cells were incubated with Nomo-1 cells (E:T = 1:1) for 72 h before sorting for RNA-seq. (**B**) UMAP visualization contrasting EGFP-NK, CD33 CAR-NK, MSLN CAR-NK, and Loop CAR-NK cells. EGFP-NK cells were used as control. (**C** and **D**) Gene Ontology (GO) analysis of differentially highly expressed genes in Loop CAR-NK cells compared with MSLN CAR-NK (C) and CD33 CAR-NK (D) cells, respectively. Each data point represented a distinct GO set. The color scale corresponded to −log_10_ transformed false discovery rate (FDR)– adjusted P values. The size of the dot was the number of genes enriched in one term. The enrichment factor was GeneRatio/BgRatio. (**E-G**) Box plots showed the expression profiles of differentially highly expressed genes related to activation and cytotoxicity of NK cells (E), participating in the RIG-1 signaling pathway (F), and involving in immunological synapse formation (G) in Loop CAR-NK cells compared with CD33 CAR-NK and MSLN CAR-NK cells.

### The fratricide of Loop CAR-NK and Loop CAR-iNK cells precludes their expansion

The expansion ability of CD33 CAR-NK cells has been impaired due to fratricide mediated by the CD33 CAR, as CD33^+^ NK cells are eliminated by CAR-expressing NK cells (*17*). Here, the expression of CD33 in UCB-derived Loop CAR-NK cells and hPSC-derived Loop CAR-iNK cells also led to fratricide. In expanded UCB-NK cells, the expression of CD33 was observed at 26.9% (**Fig. 3A)**. To assess fratricide, we sorted CD33^-^ and CD33^+^ NK cells as targets and co-incubated them with Loop CAR-NK cells (**Fig. 3**, **B** and **C)**. As expected, Loop CAR-NK cells exhibited specific cytotoxicity against CD33^+^ NK cells at the indicated effector-to-target (E:T) ratios, while no cytotoxicity was observed against CD33^-^ NK cells (**Fig. 3D).** We then sorted an equal number of EGFP-NK and Loop CAR-NK cells and cultured them consecutively without K562-mbIL-21 stimulation. Over a 12-day period, the number of Loop CAR-NK cells was lower than EGFP-NK cells, and the gap gradually widened, confirming that fratricide impaired the expansion of Loop CAR-NK cells (**Fig. 3E)**. In our previous work, we established a method for inducing iNK and CAR-iNK cells from pluripotent stem cells (*11, 35*). Here, we generated a Loop CAR-expressing hPSC line (Loop CAR-hPSCs) and then conducted the induction of Loop CAR-iNK cells (**Fig. 3F**). Briefly, Loop CAR-hPSCs were induced for two days to differentiate into Loop CAR-expressing lateral plate mesoderm cells (Loop CAR-iLPM), which were then combined with OP9 cells to form organoid aggregates. After 25 days of induction, Loop CAR-iNK cells were generated (**Fig. 3G**). The resulting Loop CAR-iNK cells expressed similar levels of CD56 and CD16 as iNK cells, with 78.6% of the cells expressing the Loop CAR (**Fig. 3H**). Moreover, Loop CAR-iNK cells showed comparable expression of typical NK cell cytotoxic effectors to the iNK cells (**fig. S3A**). Notably, the CD33 expression level in iNK cells was 69.7%, which was significantly higher than that observed in tissue-derived NK cells (**Fig. 3I**). Next, we sorted CD33^-^ and CD33^+^ iNK cells and co-incubated them with Loop CAR-iNK cells (**Fig. 3**, **J** and **K**). The Loop CAR-iNK cells showed significant cytotoxicity against CD33^+^ iNK cells compared to CD33^-^ iNK cells (**Fig. 3L**). As observed with the Loop CAR-NK cells, the number of Loop CAR-iNK cells gradually declined during a 12-day expansion period, reinforcing that fratricide significantly impaired the expansion of Loop CAR-iNK cells (**Fig. 3M**). Taken together, the expression of CD33 in Loop CAR-NK and Loop CAR-iNK cells led to fratricide, which precluded their expansion.

**Fig. 3.**
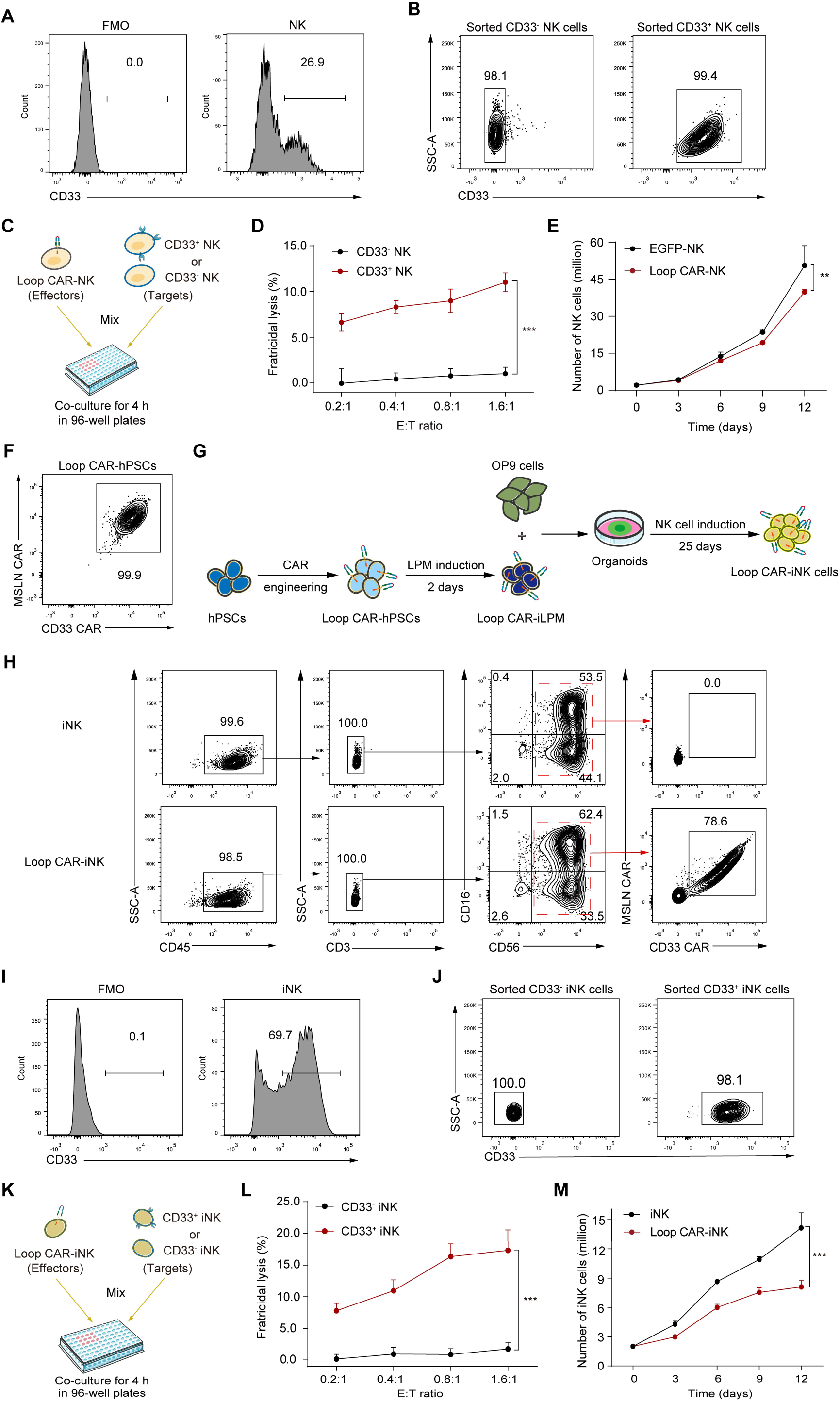
Expression of CD33 in NK or iNK cells led to fratricide and precluded expansion of Loop CAR-NK or Loop CAR-iNK cells. (**A**) Flow cytometry histograms showing the expression of CD33 in UCB-NK cells. (**B**) Representative FACS plots showing the purity of sorted CD33^-^ and CD33^+^ NK cells. (**C**) Schematic diagram of measuring the cytotoxicity of Loop CAR-NK cells against CD33^+^/CD33^-^ NK cells *in vitro*. The sorted NK cells were co-cultured with Loop CAR-NK cells for 4 h. (**D**) Statistics of the fratricidal lysis of CAR-NK cells Loop CAR-NK cells (Effectors, E) against CD33^+^ and CD33^-^ NK cells (Targets, T) at the indicated E:T ratios (*n* = 4 per group). (**E**) Statistical analysis of the cell number for EGFP-NK and Loop CAR-NK cells at indicated time points. An equal number of EGFP-NK and Loop CAR-NK cells were sorted for expansion *in vitro* (*n* = 3 per group). (**F**) Flow cytometric analysis of Loop-CAR expression in Loop CAR-hPSCs. (**G**) Schematic diagram of Loop CAR-iNK cells generated from Loop CAR-hPSCs. (**H**) Immunophenotypes (CD45^+^CD3^-^ CD56^+^CD16^+/-)^ and Loop-CAR expression (CD33 CAR^+^MSLN CAR^+^) of Loop-CAR-iNK cells. (**I**) Flow cytometry histograms showing the expression levels of CD33 in iNK cells derived from hPSCs. (**J**) Representative FACS plots showing the the purity of sorted CD33^-^ and CD33^+^ iNK cells. (**K**) Schematic diagram of measuring the cytotoxicity of Loop CAR-iNK cells against CD33^+^ and CD33^-^ iNK cells *in vitro*. (**L**) Statistics of fratricidal lysis of Loop CAR-iNK cells (Effectors, E) against CD33^+^ and CD33^-^ iNK cells (Targets, T) at indicated E:T ratios (*n* = 4 per group). (**M**) Statistical analysis of the cell number for iNK and Loop CAR-iNK cells at indicated time points (*n* = 3 per group). An equal number of iNK and Loop CAR-iNK cells were started for culture. Data represent the mean ± SEM. Statistics analysis: one-way ANOVA test (D, L) and two-way ANOVA test (E, M). NS: not significant; \*\**P* < 0.01, \*\*\**P* < 0.001.

### CD33 inactivation in iNK cells preserves their broad-spectrum cytotoxicity and facilitates the expansion of Loop CAR-iNK cells

To address the fratricide caused by CD33 expression in Loop CAR-iNK cells, we generated a CD33 gene knockout hPSC (CD33^KO^-hPSC) line using the CRISPR/Cas12i system. Vectors carrying Cas12i, sgRNA1 (5’-TTCAGAAGTGGCCGCAAGGGAAG-3’), sgRNA2 (5’-TTGCTAGTCATCCCCAAATATCC-3’), and a neomycin resistance gene were transduced into hPSCs (**Fig. 4A**). Following neomycin selection, single clones were isolated and genotyped using four pairs of PCR primers (**fig. S4**, **A** and **B**). Ultimately, we identified a CD33^KO^-hPSC clone with the deletion of exon 3 and exon 4 in one allele of CD33 and the insertion of a loxp-NeoR-loxp sequence in the other (**Fig. 4B** and **fig. S4**, **C** and **D**). Subsequently, CD33^KO^-hPSCs successfully generated CD33-deficient iNK cells (CD33^KO^-iNK) with immunophenotype similar to iNK cells (**Fig. 4C**). Moreover, the yields of CD33^KO^-iNK cells were comparable to those of iNK cells (**Fig. 4D**). Importantly, CD33^KO^-iNK cells exhibited cytotoxicity against K562 cells equivalent to that of iNK cells, demonstrating that the loss of CD33 did not compromise their broad-spectrum cytotoxicity (**Fig. 4E**). We then introduced the Loop CAR into CD33^KO^-hPSCs to establish the CD33^KO^-Loop CAR-hPSC line. As anticipated, these cells successfully differentiated into CD33^KO^-Loop CAR-iNK cells, which exhibited expression levels of CD56 and CD16 comparable to those of iNK cells, with a CAR expression rate of 89.3% (**Fig. 4**, **F** and **G**). Furthermore, the expression of typical activating and inhibitory receptors, as well as cytotoxic granules, was consistent with that of iNK cells (**fig. S4E**). Most notably, CD33^KO^-Loop CAR-iNK cells demonstrated a similar expansion capacity to that of iNK cells, indicating that CD33 disruption effectively resolved the fratricide issue and restored the proliferation of Loop CAR-iNK cells **(Fig. 4H)**. In conclusion, we successfully mitigated the fratricide issue in Loop CAR-iNK cells by knocking out CD33 in hPSCs, thereby preserving their broad-spectrum cytotoxicity and restoring their expansion.

**Fig. 4.**
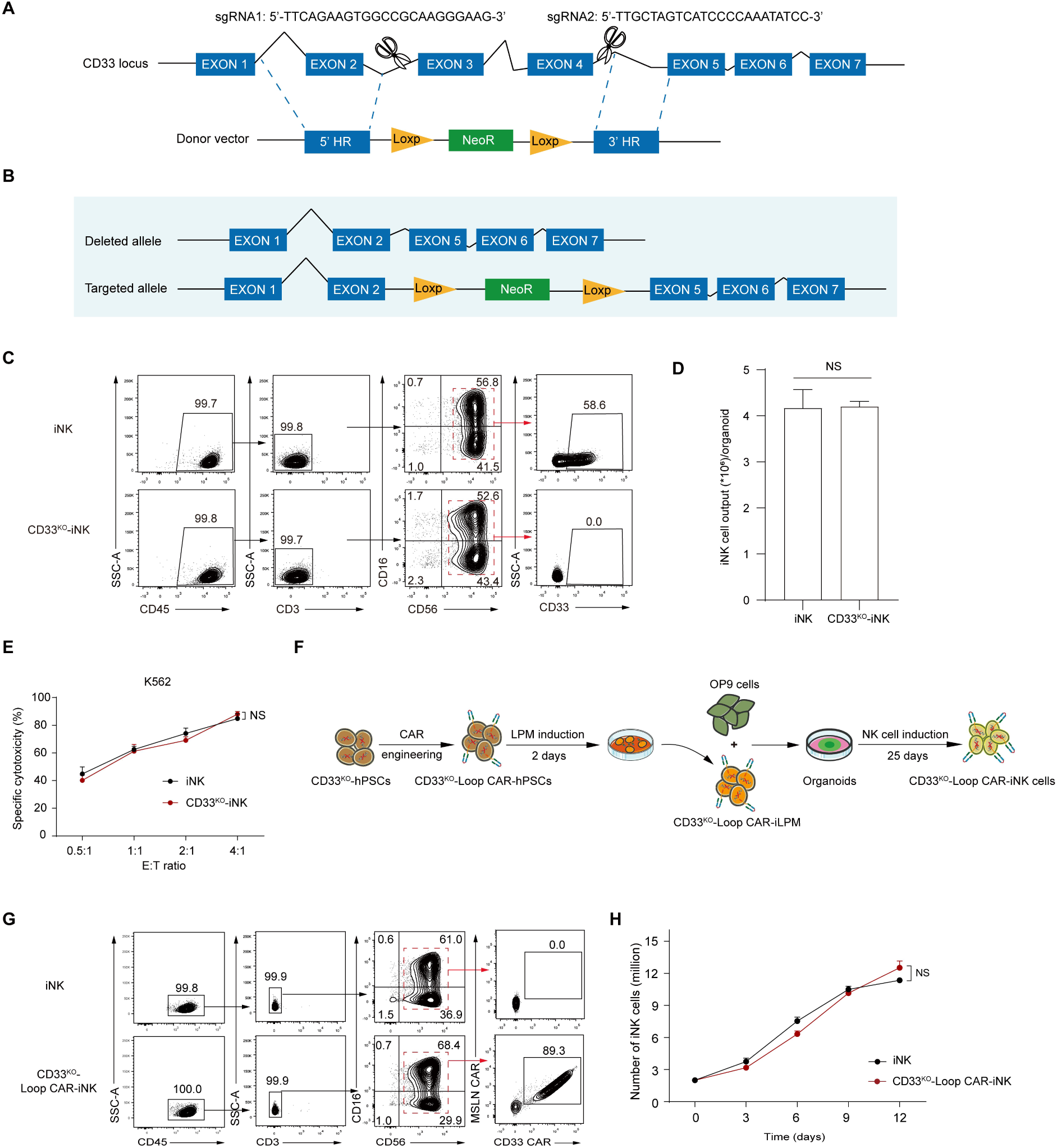
Inactivation of CD33 restored the expansion of Loop CAR-iNK cells. (**A**) Schematic diagram of CRISPR/Cas12i-targeted CD33 gene strategy. sgRNA: single guide RNA; HR: homologous region; NeoR: neomycin resistance gene. (**B**) Schematic diagram showing the genotype of the established CD33^KO^-hPSCs. (**C**) Flow cytometric analysis of iNK and CD33^KO^-iNK cells. (**D**) Statistics of the yields of iNK and CD33^KO^-iNK cell (Day 27) induced from hPSCs and CD33^KO^-hPSCs (*n* = 3 repeats). (**E**) Statistics of cytotoxicity of iNK and CD33^KO^-iNK cells (Effectors, E) against K562 cells (Targets, T) at the indicated E:T ratios (*n* = 4 per group). (**F**) Schematic diagram of CD33^KO^-Loop CAR-iNK cells induced from CD33^KO^-Loop CAR-hPSCs. (**G**) Immunophenotypes (CD45^+^CD3^-^CD56^+^CD16^+/-^) and Loop-CAR expression in iNK and CD33^KO^-Loop CAR-iNK cells. (**H**) Statistical analysis of the cell number for iNK and CD33^KO^-Loop CAR-iNK cells at indicated time points. An equal number of iNK and Loop CAR-iNK cells were started for culture (*n* = 3 repeats). Data represent the mean ± SEM. Statistics analysis: two-tailed Student’s t test (D), one-way ANOVA test (E) and two-way ANOVA test (H). NS: not significant.

### Genetic disruption of CD33 in Loop CAR-iNK cells enhances their specific cytotoxicity

To evaluate the specific cytotoxicity of CD33^KO^-Loop CAR-iNK cells, we incubated them with Nomo-1 cells or MSLN-expressing HL-60 cells (MSLN^+^ HL-60) (**Fig. 5**, **A** and **B**). Notably, CD33^KO^-Loop CAR-iNK cells exhibited significantly enhanced cytotoxicity compared to Loop CAR-iNK cells, suggesting that fratricide in the latter compromised their specific cytotoxicity (**Fig. 5**, **C** and **D**). To further evaluate the durability of the cytotoxic response, we performed serial killing assays by analyzing the cytotoxicity of CD33^KO^-Loop CAR-iNK cells (**Fig. 5E**). Compared to Loop CAR-iNK cells, CD33^KO^-Loop CAR-iNK cells consistently demonstrated superior specific cytotoxicity against Nomo-1 cells in each round killing assay (**Fig. 5F**). Additionally, we assessed the expression levels of key effector molecules, including TNF-α, IFN-γ, and CD107a, in CD33^KO^-Loop CAR-iNK cells following incubation with Nomo-1 and MSLN^+^ HL-60 cells. The results revealed that CD33^KO^-Loop CAR-iNK cells expressed significantly higher levels of these molecules compared to Loop CAR-iNK cells, further corroborating their enhanced effector function (**Fig. 5**, **G-O**). Collectively, these findings underscore the critical role of CD33 disruption in restoring and strengthening the specific cytotoxicity of Loop CAR-iNK cells, demonstrating its importance in overcoming fratricide-induced functional impairments.

**Fig. 5.**
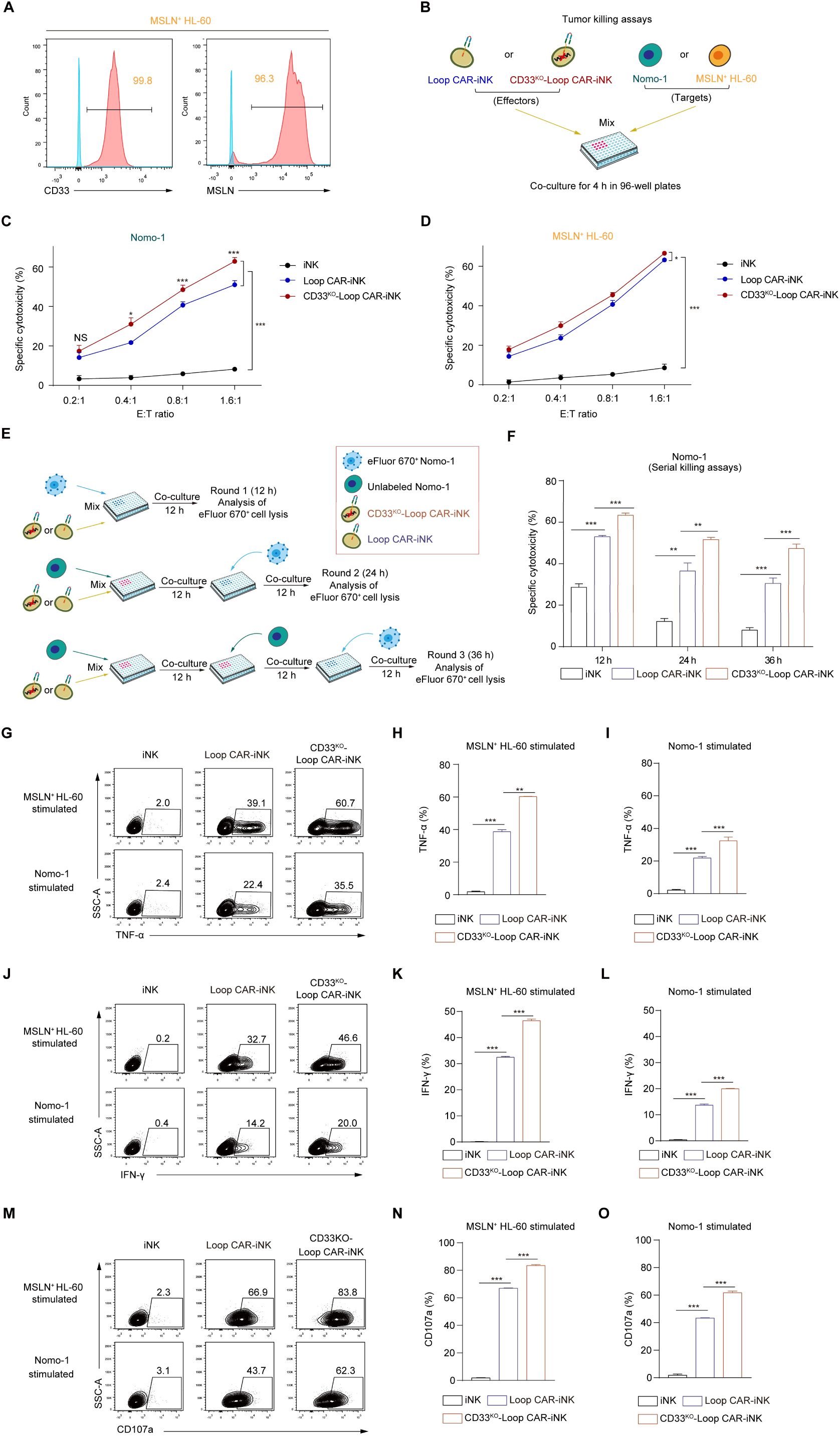
CD33^KO^-Loop CAR-iNK cells exhibited superior tumor-killing abilities than Loop CAR-iNK cells. (**A**) Flow cytometry histograms showing the expression of CD33 and MSLN in HL-60 cells engineered with MSLN (MSLN^+^HL-60). (**B**) Schematic diagram showing Nomo-1 or MSLN^+^ HL-60 cells killed by Loop CAR-iNK or CD33^KO^-Loop-CAR-iNK cells *in vitro*. (**C** and **D**) Statistics of specific cytotoxicities analysis of iNK, Loop CAR-iNK and CD33^KO^-Loop CAR-iNK cells (Effectors, E) against Nomo-1 (C) and CD33^+^MSLN^+^ HL-60 cells (D) (Targets, T) at the indicated E:T ratios (*n* = 4 per group). (**E**) Schematic diagram of the serial killing assays of iNK, Loop CAR-iNK, and CD33^KO^-Loop CAR-iNK cells against Nomo-1 cells. Each group of iNK cells was co-cultured with Nomo-1 cells for 12 h per round at an E:T ratio of 1:1. The fresh Nomo-1 cells were added to the NK cell residues and incubated for every other 12 h. The lysis of Cell Proliferation Dye eFluor 670-labeled (eFluor 670^+^) Nomo-1 cells in each round represented the cytotoxicity. (**F**) Statistics of specific cytotoxicity of iNK, Loop CAR-iNK, and CD33^KO^-Loop CAR-iNK cells in serial killing assays (*n* = 4 per group). (**G**, **J**, **M**) Representative flow cytometry plots showing the expression of TNF-α (G), IFN-γ (J), and CD107a (M) in each group of iNK cells stimulated by MLSN^+^ HL-60 and Nomo-1 cells. (**H** and **I**) Statistical analysis of the expression levels of TNF-α (*n* = 3 repeats). (**K** and **L**) Statistical analysis of the expression levels of IFN-γ (*n* = 3 repeats). (**N** and **O**) Statistical analysis of the expression levels of CD107a (*n* = 3 repeats). Data represent the mean ± SEM. Statistics analysis: one-way ANOVA test. NS: not significant; \**P* < 0.05, \*\**P* < 0.01, \*\*\**P* < 0.001.

### CD33^KO^-Loop CAR-iNK cells effectively suppress AML tumor development in xenograft animals

To further evaluate the anti-tumor efficacy of CD33^KO^-Loop CAR-iNK cells, we employed an AML xenograft mouse model, using iNK cells as control effectors (**Fig. 6A**). MSLN^+^ HL-60 cells were intravenously injected into NCG mice (5 × 10^5^ cells/mouse) on day -1 to establish the AML xenograft models. Mice with comparable tumor burdens were randomly assigned to three groups (Tumor only, iNK, and CD33^KO^-Loop CAR-iNK). CD33^KO^-Loop CAR-iNK cells were administered intravenously at a dose of 1 × 10⁷ cells/mouse on days 0 and 3 (**Fig. 6B**). Tumor progression was monitored weekly using bioluminescence imaging (BLI). The results showed that the tumor burden in mice treated with CD33^KO^-Loop CAR-iNK cells was significantly lower than in those treated with iNK cells, demonstrating the potent anti-tumor activity of CD33^KO^-Loop CAR-iNK cells (**Fig. 6**, **C** and **D**). Moreover, the enhanced tumor suppression translated into a significant survival benefit, with CD33^KO^-Loop CAR-iNK-treated mice achieving a median survival of 42 days, compared to shorter survival in the control groups (**Fig. 6E**). Collectively, these findings demonstrate that CD33^KO^-Loop CAR-iNK cells effectively eliminate AML tumor cells *in vivo* and prolong the survival of xenograft models, underscoring their therapeutic potential for clinical application.

**Fig. 6.**
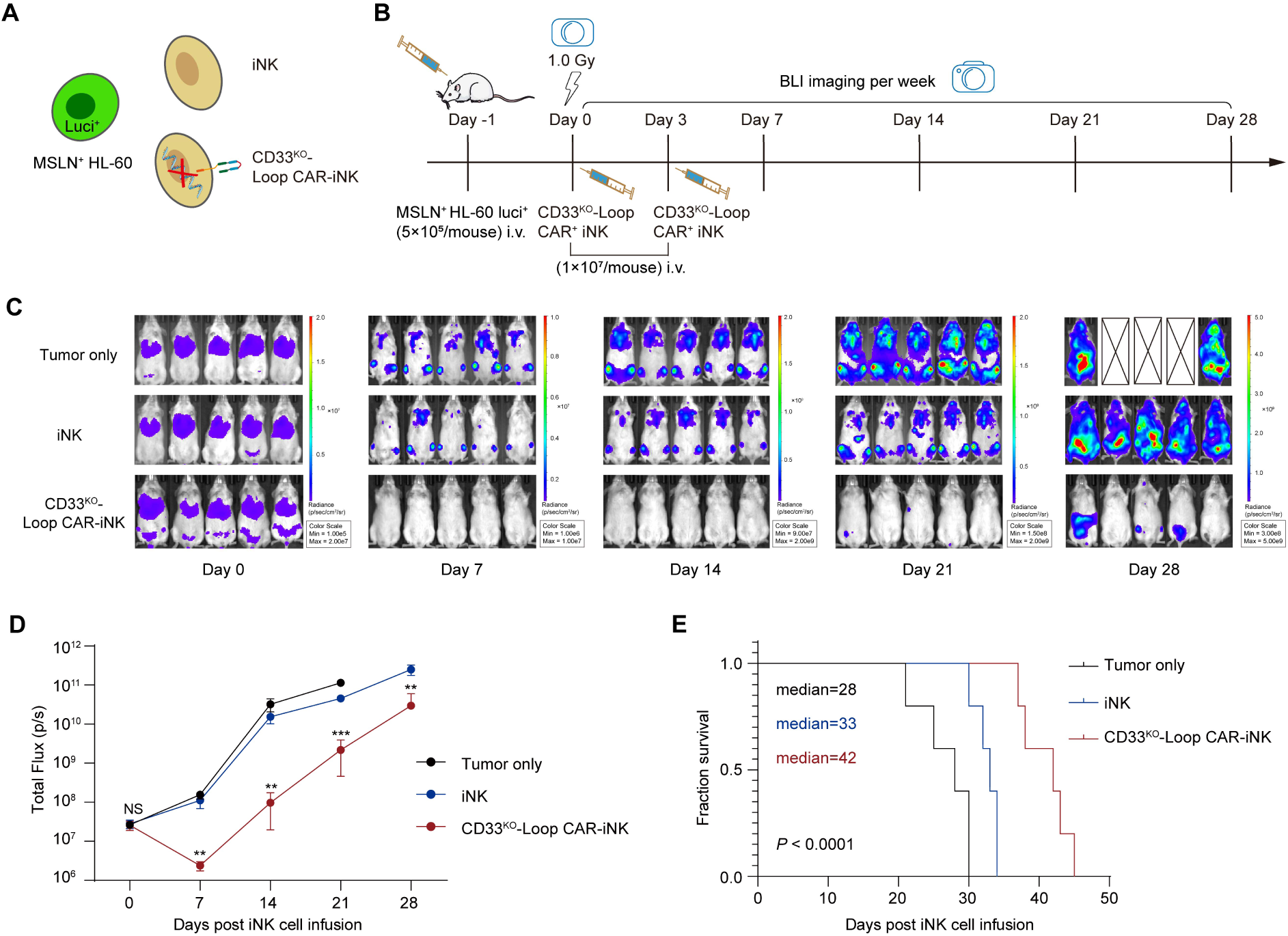
CD33^KO^-Loop CAR-iNK cells efficiently suppressed tumor progression *in vivo*. (**A**) Schematic diagram of the tumor cells and therapeutic cells used for *in vivo* assay. (**B**) Schematic diagram of *in vivo* studies. Luciferase-expressing (luci^+^) MSLN^+^ HL-60 cells were used to generate mouse xenograft models (5 × 10^5^ cells/mouse, i.v.). Two doses of CD33^KO^-Loop CAR-iNK cells (1× 10^7^ cells/mouse per dose) were injected into tumor-bearing mice for therapy. (**C**) Bioluminescence imaging (BLI) of xenograft models of each group (*n* = 5 per group). (**D**) Statistics of each group’s total flux (p/s) (CD33^KO^-Loop CAR-iNK vs. iNK). Statistics analysis: two-tailed Student’s *t*-test. NS: not significant; \*\**P* < 0.01, \*\*\**P* < 0.001. (**E**) Kaplan-Meier survival curves of the experimental groups. Statistics analysis: two-tailed log-rank test. Tumor only vs. iNK, \*\**P* < 0.01, CD33^KO^-Loop CAR-iNK vs. iNK, \*\**P* < 0.01.

## DISCUSSION

The dual CAR strategy is an effective approach to prevent tumor escape caused by antigen heterogeneity or loss. Bivalent CAR, one CAR containing two scFvs, which can result in higher transduction efficiency than bicistronic CAR, two single-targeted CARs. Bivalent CAR can be categorized into tandem CAR and loop CAR (*36*). Previous studies have reported that loop CAR exhibited superior anti-tumor potency than tandem CAR (*37, 38*). In this study, we developed a bivalent loop CAR targeting CD33 and mesothelin (MSLN) (CD33-MSLN loop CAR, denoted as Loop CAR). When the Loop CAR expression cassette was introduced into umbilical cord blood NK (UCB-NK) cells, the resulting Loop CAR-NK cells exhibited superior cytotoxicity against CD33 and MSLN-expressing tumor cells, including Nomo-1 and primary CD33^+^MSLN^+^ AML cells. Furthermore, Loop CAR-NK cells demonstrated comparable cytotoxicity against CD33^+^MSLN^−^ and CD33^−^MSLN^+^ tumor cells compared to CD33 CAR-NK and MSLN CAR-NK cells. The accumulation of Loop CAR proteins at the immunological synapse between Loop CAR-NK cells and Nomo-1 cells likely contributes to their enhanced specific cytotoxicity (*39, 40*). Additionally, Nomo-1 cells stimulated Loop CAR-NK cells to express higher levels of CD107a, a membrane protein associated with NK cell cytotoxic activity (*41*). Transcriptome analysis revealed that Loop CAR-NK cells exhibited upregulated signaling pathways related to cell division and activation after incubation with Nomo-1 cells. Genes such as *NCAM1*, *PTK2B*, and *DOCK8*, which are involved in immunological synapse formation and cytotoxic function, were significantly upregulated (*31–34, 42*).

Consistent with the previous report (*23*), a fraction of UCB-NK cells naturally express CD33. Our findings confirmed that hPSC-derived iNK cells exhibit even higher levels of CD33 expression compared to UCB-NK cells. This led to fratricide in both Loop CAR-NK and Loop CAR-iNK cells, ultimately impairing their expansion. Similar fratricide-induced expansion impairments have been previously reported in human tissue-derived CD33 CAR-NK cells (*17*), highlighting the necessity of addressing this challenge. To overcome the fratricide issue, we genetically disrupted CD33 in hPSCs (CD33^KO^-hPSCs) using CRISPR/Cas technology. Previous studies have shown that CD33 knockout in hematopoietic stem cells allows for the development of a functional hematopoietic system without impairing differentiation potential (*21*). Our results demonstrated that CD33^KO^-hPSCs successfully differentiated into iNK cells without affecting their induction efficiency or anti-tumor activity. Although CD33 has been considered an inhibitory receptor due to its cytoplasmic tyrosine-based inhibitory motifs (ITIMs) (*43*). We observed no significant differences in cytotoxicity against K562 cells between CD33^KO^-iNK and iNK cells.

As expected, CD33^KO^-Loop CAR-hPSCs efficiently generated CD33^KO^-Loop CAR-iNK cells. The elimination of CD33 expression successfully prevented fratricide and restored the expansion capacity of Loop CAR-iNK cells. Interestingly, CD33^KO^-Loop CAR-iNK cells demonstrated superior specific cytotoxicity against Nomo-1 and MSLN^+^ HL-60 cells compared to Loop CAR-iNK cells, indicating that fratricide in the latter not only impaired their expansion but also reduced their specific cytotoxicity. Moreover, CD33^KO^-Loop CAR-iNK cells significantly suppressed tumor progression and prolonged the survival of AML xenograft mice. We used MSLN^+^ HL-60 cells, an M2 subtype of acute myeloid leukemia (AML), to construct the tumor models, as these cells exhibit high tumorigenic potential *in vivo*. Additionally, we attempted to use Nomo-1 cells, an M5a subtype of acute monocytic leukemia, to establish a tumor model. However, the tumorigenic efficiency was suboptimal with Nomo-1 cells. Given NK cells’ relatively lower genome editing efficiency compared to T cells (*44*), targeting endogenous CD33 at the pluripotent stem cell (PSC) stage represents a more practical approach to avoid potential fratricide. With their amenability to genetic engineering, clonal selection capabilities, and lack of T cell contamination, PSC-derived NK cells hold promise as standardized, off-the-shelf immunotherapy products (*45, 46*).

Despite significant progress in identifying AML-specific antigens as therapeutic targets, challenges remain due to their genomic and phenotypic heterogeneity and overlapping expression with normal hematopoietic cells (*5*). CD33, expressed in over 80% of AML cells, is absent from primitive stem cells and multipotent progenitor cells, making it an attractive target for AML immunotherapy (*47, 48*). Recent clinical data have demonstrated the safety and efficacy of functionally enhanced CD33 CAR-NK cells (*10*). MSLN, a well-established target in solid tumors, is expressed in approximately 36% of pediatric AML cases and 14% of adult AML cases, but is virtually absent in normal hematopoietic cells (*4*). MSLN CAR-T cells can efficiently eliminate MSLN-positive (MSLN^+^) AML cells in xenograft models derived from cell lines and patients (*5*). However, MSLN is actively cleaved from the cell membrane and is not highly expressed in AML cells. Strategies such as small molecules (e.g., INCB7839, GM6001) and immunotoxins (e.g., RG7787, SS1P) have been developed to upregulate MSLN expression and reduce shedding, thereby enhancing its viability as a therapeutic target (*5, 49, 50*). Our findings demonstrated that CD33-MSLN loop CAR-NK cells exhibited superior cytotoxicity against primary AML cells expressing both CD33 and MSLN (23.5%-57.8%). Moreover, CD33^KO^-CD33-MSLN loop CAR-iNK cells displayed robust anti-tumor efficacy both *in vitro* and in xenograft models. These results indicate that the CD33-MSLN loop CAR design effectively enhances the tumor-killing potential of NK cells and supports the successful development of a dual-targeting CAR strategy. Additionally, by genetically inactivating CD33, we effectively addressed the fratricide issue, further improving the function of CD33-MSLN loop CAR-iNK cells. In conclusion, our study provides compelling preclinical evidence supporting the further evaluation of CD33-MSLN loop CAR-NK and CD33^KO^-CD33-MSLN loop CAR-iNK cells as promising candidates for clinical trials in the treatment of MSLN^+^ AML.

## MATERIALS AND METHODS

### Study design

The objective of this study was to develop a more efficient therapeutic CAR-NK cell for the treatment of acute myeloid leukemia (AML). This was achieved by designing a bivalent CD33-MSLN loop CAR (Loop CAR) and inactivating CD33, thereby generating Loop CAR-iNK cells that avoid fratricide and effectively target and kill MSLN^+^CD33^+^ AML cells. The construction of Loop CAR was described in the results. The functionality of the Loop CAR was validated by cloning its expression elements into the SFG vector and employing the RD114 retroviral system in conjunction with umbilical cord blood NK cells to generate Loop CAR-NK cells. The superior *in vitro* cytotoxicity of Loop CAR-NK cells compared to CD33 CAR-NK and MSLN CAR-NK cells was assessed through tumor killing assays against CD33^+^MSLN^+^ tumor cell lines (Nomo-1 and MSLN^+^ HL-60). Furthermore, the expression of CD107a in CAR-NK cells stimulated by Nomo-1 cells was analyzed to evaluate degranulation capacity. Single-cell RNA sequencing (scRNA-seq) using the 10× Genomics platform further demonstrated the enhanced effector functions of loop CAR-NK cells at the transcriptomic level following co-incubation with Nomo-1 cells.

To extend this strategy, the Loop CAR was introduced into hPSCs to generate Loop CAR-iNK cells. However, functional assays revealed that both Loop CAR-NK and Loop CAR-NK iNK cells exhibited fratricide by targeting CD33^+^ NK cells or iNK cells, leading to limited expansion. To address this challenge, the CRISPR/Cas12i system was employed to generate a CD33 knockout (CD33^KO^) hPSC line. We first confirmed that CD33 disruption did not affect the immunophenotype or cytotoxic function of iNK cells through flow cytometric immunophenotypic analysis and tumor-killing assays against K562 cells. Subsequently, the Loop CAR was introduced into the CD33^KO^-hPSCs to produce CD33^KO^-Loop CAR-iNK cells. Flow cytometric analysis and *in vitro* expansion experiments confirmed that the loss of CD33 did not alter the immunophenotypic characteristics of Loop CAR-iNK cells but significantly enhanced their expansion potential. To validate the improved *in vitro* effector functions of CD33^KO^-Loop CAR-iNK cells, tumor-killing assays and flow cytometry analysis of CD107a expression and effector molecule secretion (IFN-γ and TNF-α) were conducted following stimulation with Nomo-1 and MSLN^+^ HL-60 cells. To assess the *in vivo* anti-tumor efficacy of CD33^KO^-Loop CAR-iNK cells, AML xenograft models were established using MSLN^+^ HL-60 cells in NCG mice. Tumor-bearing mice were treated with either iNK or CD33^KO^-Loop CAR-iNK cells, and therapeutic efficacy was monitored through weekly bioluminescence imaging. Blinding was not employed in the experiments. Each experiment was conducted independently on different days using different batches of reagents. Experimental data were repeated to ensure an adequate number of biological replicates. Details regarding sample sizes, replicates, and statistical analyses are provided in the figures, figure legends, and supplementary materials.

### Cell culture

Human embryonic kidney (HEK) 293T cells (ATCC) were cultured in Dulbecco’s Modified Eagle Medium (DMEM, Gibco) supplemented with 10% fetal bovine serum (FBS, HUANGKE). Human cervical cancer HeLa cells (Procell Life Science & Technology Co., Ltd) were maintained in Minimum Essential Medium α (MEMα, Gibco) with 10% FBS. Mesothelin-expressing (MSLN^+^) AML cell lines, Nomo-1 (COBIOER), and luciferase-expressing Nomo-1 were maintained in RPMI 1640 (Gibco) supplemented with 10% FBS. Human gastric adenocarcinoma AGS cells (Procell Life Science & Technology Co., Ltd) were cultured in Ham’s F12 (Gibco) supplemented with 10% FBS. HL-60 (Procell Life Science & Technology Co., Ltd) and luciferase-expressing MSLN^+^ HL-60 MSLN^+^ HL-60 luci^+^ cells were cultured in Iscove’s Modified Dulbecco’s Medium (IMDM, Gibco) supplemented with 20% FBS. To generate MSLN-expressing HL-60 cell line, the cDNA encoding MSLN was inserted into the PiggyBac expression vector (SBI) to generate a recombinant vector. The MSLN-expressing PiggyBac vector and the transposase expression vector were electroporated into HL-60 cells using Electroporator EX+ (Celetrix). Seven days after electroporation, HL-60 cells were sorted for two rounds to establish the MSLN-expressing HL-60 cell line. The PSC line (Q380-ESC) was provided by the National Stem Cell Resource Center, Institute of Zoology, Chinese Academy of Sciences. All hPSCs were kept in Essential 8 Medium (Gibco) on Vitronectin-Recombinant Human Protein (rhVTN-N, Gibco) coated plates. The hPSCs differentiation toward iNK and the related anti-tumor activity assessments of iNK cells in animals were approved by the Biomedical Research Ethics Committee of the Institute of Zoology, Chinese Academy of Sciences.

### Generation of CAR constructs and CAR-modified NK cells

The monospecific CAR constructs included a signal peptide (SP), the anti-CD33 or anti-mesothelin (MSLN) single-chain variable fragment (scFv) (*51, 52*), the CD8α hinge domain, a transmembrane domain (TMD) of CD8, and a signaling domain (SD) of CD3ζ. The Loop CAR construct was designed using the same backbone, with the scFv arrangement as follows: CD33 (V_L)_-MSLN (V_H_)-MSLN (V_L_)-CD33 (V_H_) were fused with the G4S linker and the whitlow linker (*53*). The EGFP coding sequence or CAR constructs were cloned into SFG recombinant retroviral vectors (*54*). Retroviruses carrying the CAR vectors were generated as previously described (*25*). Umbilical cord blood (UCB) units were provided by Guangdong Cord Blood Bank (Guangzhou, China). CD3^-^ umbilical cord blood mononuclear cells (UCBMCs) were isolated, and activated NK cells were produced as previously described (*25*). On day 0, activated NK cells were transduced with retroviral supernatants at a multiplicity of infection (MOI) of 5, following the Vectofusin-1 (Miltenyi Biotec) based transduction protocol. Infection rates were assessed by flow cytometry on day 2. CAR-NK cells were harvested for use after 4 days of expansion. The CAR-NK cells were sorted and expanded for 12 days without stimulation by K562-mbIL-21 cells (Hangzhou Zhongying Biomedical Technology Co., Ltd).

### Flow cytometry

Transduced UCB-NK cells were stained with fluorochrome-conjugated antibodies against CD3 (UCHT1), CD56 (HCD56) (Biolegend), CD33 scFv, and MSLN scFv (ACROBiosystems) for analysis of CAR expression using a BD LSRFortessa flow cytometer (BD Biosciences). Anti-CD56, anti-CD33 scFv, and anti-MSLN scFv antibodies were used to sort CAR-NK cells using Sony MA900 (SONY). Human anti-CD33 antibody (P67.6, Biolegend) and human anti-MSLN antibody (Abcam) were used to determine the expression levels of CD33 and MSLN in tumor cell lines and primary AML cells. The immunophenotypes of iNK cells were analyzed using antibodies against CD45 (HI30, Biolegend), CD3, CD56, CD16 (3G8, Biolegend), CD33 scFv and MSLN scFv. To evaluate the expression of NK receptors and effector molecules, we used antibodies against CD56, NKp44 (P44-8), NKp30 (P30-15), NKG2D (1D11), CD69 (FN50), CD16, CD319 (162.1), CD226 (11A8), CD96 (NK92.39), CD94 (HP-3D9), NKG2A (S19004C), Perforin (dG9), GzmB (QA18A28) (all from Biolegend), CD33 scFv and MSLN scFv. Cells were resuspended in DAPI (Sigma-Aldrich) or Dulbecco’s phosphate-buffered saline (BasalMedia Technologies) for analysis with the BD LSRFortessa cytometer (BD Biosciences). Data were analyzed with FlowJo software (FlowJo_v10.8.1).

### Cytotoxicity assays

For cytotoxicity assessment, target cells, labeled with eBioscience^TM^ Cell Proliferation Dye eFluor^TM^ 670 (eFluor 670) (Invitrogen), were co-cultured with NK cells in the indicated effector (E) to target (T) ratios (E:T). Target cell death was quantified using the BD LSRFortessa flow cytometer or BD LSRFortessa X-20 cytometer (BD Biosciences) by measuring the percentage of DAPI^+^ among eFluor 670-labeled cells. In serial killing assays, iNK cells received three rounds tumor challenge. Round 1: eFluor 670-labeled (eFluor 670^+^) Nomo-1 cells were mixed with relevant iNK cells and co-incubated for 12 h (E:T = 1:1). Analysis of the cell lysis of eFluor 670^+^ Nomo-1 cells at 12 h represents the Round 1 cytotoxicity of the relevant iNK cells. Round 2: Unlabeled Nomo-1 cells were mixed with relevant iNK cells and co-incubated for 12 h (E:T = 1:1). Equal numbers of eFluor 670-labeled Nomo-1 cells were added and co-incubated for the next 12 h. Analysis of the cell lysis of eFluor 670^+^ Nomo-1 cells at 24 h represents the Round 2 cytotoxicity of the relevant iNK cells. Round 3: Unlabeled Nomo-1 cells were mixed with relevant iNK cells (E:T = 1:1). Equal numbers of unlabeled Nomo-1 cells and eFluor 670-labeled Nomo-1 cells were added at 12 h and 24 h respectively. Analysis of the cell lysis of eFluor 670^+^ Nomo-1 cells at 36 h represents the Round 3 cytotoxicity of the relevant iNK cells. Specific cytotoxicity was calculated using the formula: (percentage of specific death – percentage of spontaneous death) / (1 – percentage of spontaneous death) × 100%.

### Assessment of CD107a, TNF-α, and IFN-γ by flow cytometry

To evaluate CD107a expression, NK or iNK cells were co-cultured with or without Nomo-1 or MSLN^+^ HL-60 cells at an E:T ratio of 0.5:1 for 4 h. After incubation, cells were stained with antibodies against CD56, CD107a (H4A3, Biolegend), CD33 scFv, and MSLN scFv. To evaluate TNF-α and IFN-γ expression, iNK cells were co-cultured with or without Nomo-1 or MSLN^+^ HL-60 cells at an E:T ratio of 0.5:1 for 2 h, followed by adding BFA/Monensin (MULTISCIENCES) for additional 2-h incubation. After incubation, cells were stained with antibodies against CD56, CD33 scFv, and MSLN scFv. FIX &PERM Kit (MULTISCIENCES) was used for fixation and permeabilization, followed by intracellular staining for TNF-α (Biolegend, MAb11) or IFN-γ (Biolegend, 4S.B3). The cells were analyzed with BD LSRFortessa X-20 cytometer (BD Biosciences). Data were analyzed with FlowJo software (FlowJo_v10.8.1).

### Immunofluorescence imaging

Immunofluorescence imaging of immune synapses of Loop CAR-NK cells by confocal microscopy as previously described (*55*). Briefly, immune synapses of Loop CAR-NK cells were formed by suspending NK cells at 5 × 10^6^/mL and Nomo-1 target cells at 2.5 × 10^6^/mL. Equal volumes of Loop CAR-NK cell and Nomo-1 target cell suspensions were mixed and incubated to form conjugates. Loop CAR-NK and Nomo-1 cell conjugates were permeabilized using the FIX & PERM kit (MULTISCIENCES) and stained with antibodies against phalloidin (Thermo Fisher Scientific), CD33 scFv, MSLN scFv, pericentrin (Abcam), and perforin (Biolegend). Sequential scanning with a Leica TCS SP8 laser scanning confocal microscope imaged NK cell immune synapses. The images were acquired using LASAF software (Leica).

### Quantitative real-time PCR

The *MSLN* transcripts were reverse transcribed into cDNA using HiScript III All-in-One RT SuperMix Perfect for qPCR (Vazyme) and quantified by quantitative real-time PCR using Hieff® qPCR SYBR® Green Master Mix (Yeasen Biotechnology). *GAPDH* was used for normalization. The relative gene expression levels were calculated as 2^-ΔCt^. All primers used in this study were listed in **table S1**.

### Primary samples

Primary AML cells were isolated from AML diagnostic bone marrow or peripheral blood samples obtained from the State Key Laboratory of Experimental Hematology, National Clinical Research Center for Blood Diseases, Institute of Hematology & Blood Diseases Hospital, Chinese Academy of Medical Science & Peking Union Medical College, and Department of Hematology, Guangdong Provincial People’s Hospital. Primary AML cells were thawed for immunophenotype analysis of MSLN expression and cultured in IMDM supplemented with 20% FBS, 10 ng/ml GM-CSF (MedChemExpress), 100 U/ml DNase I (Sigma Aldrich), and 50 uM GM6001 (Selleck) (a matrix metalloprotease inhibitor that stabilizes MSLN expression on the cell surface (*5*) for cytotoxicity assays. *In vitro* killing assays were performed as mentioned above. The use of patient samples was carried out by the provisions of the Declaration of Helsinki. All patient samples were collected with patient consent signatures and reviewed and approved by the Ethics Committee of the Hospital for Hematology & Blood Diseases and the Guangdong Provincial People’s Hospital, respectively.

### Single-cell RNA-sequencing and data analysis

FACS-sorted EGFP-NK (CD56^+^EGFP^+^) and CAR-NK cells (CD56^+^CAR^+^) were activated with eFluor 670 labeled (eFluor 670^+^) Nomo-1 cells for 72 h. Fifty thousand cells sorted (eFluor 670^-^ and CD56^+^) from activated NK populations underwent RNA-sequencing (10× Genomics). Droplet-based scRNA-seq datasets were generated using a Chromium system (10× Genomics, PN120263) following the manufacturer’s instructions. The raw data were uploaded to the public database of Genome Sequence Archive (HRA008163). Subsequently, these data sets were aligned and quantified using the CellRanger software package (version 6.0.0) and subjected to Seurat (version 3.2.3) for further analysis. To pass quality control, cells were required to have at least 200 genes detected, less than 7,500 genes detected, and less than 20% of aligned reads mapped to mitochondrial genes. 15,408, 17269, 15273, and 16734 cells from EGFP-NK, CD33 CAR-NK, MSLN CAR-NK, and Loop CAR-NK scRNA-seq datasets that passed quality control were merged and used for downstream analysis. The Unique Molecular Identifier (UMI) value was normalized using Seurat’s ‘LogNormalize’ and ‘ScaleData’ methods. The Principal Component Analysis (PCA) used 2,000 highly variable genes (as defined by Seurat), and clustering was performed on the most variable principal components (PCs). The most variable PCs were determined by the Jackstraw method. Then, the 36 most variable PCs were used for clustering and Uniform Manifold Approximation and Projection (UMAP) analysis with resolution 0.2 and other default parameters. The median and dot plots for gene expression were plotted using the Seurat VlnPlot function and the ggplot2 package. Differential expression genes (DEGs) were identified using the neurinoma test with the parameter ‘min.pct = 0.25, logfc threshold = 0.25’. Gene ontology (GO) enrichment analysis for DEGs was performed by the clusterProfiler package (version 4.2.2) with BH adjusted cutoff value = 0.05. Box plots of the normalized expression of DEGs were plotted using the ggplot2 package. Volcano plots were plotted using ggplot2 and the ggrepel package.

### Regeneration of CAR-iNK cells

CAR-expressing hPSCs were engineered using the piggyBac transposon system, as mentioned above. CAR-iNK cells were regenerated from CAR-transfected hPSCs and harvested for use as previously described (*11*). Briefly, CAR-hPSCs were performed a two-day induction to produce CAR lateral plate mesoderm cells (CAR-iLPM). Then 2 × 10^4^ CAR-iLPM were combined with 5 × 10^5^ OP9 cells to form organoid aggregates and seeded in the transwell (Corning, 3450). Subsequently, the medium was changed every 2-3 days to induce NK cell regeneration. After 25 days of induction, mature CAR-iNK cells were harvested for use.

### CRISPR/Cas12i-mediated CD33 knockout in hPSCs

Two sgRNA sequences, sgRNA1 (TTCAGAAGTGGCCGCAAGGGAAG) and sgRNA2 (TTGCTAGTCATCCCCAAATATCC), targeting exon 3 and exon 4 of CD33 respectively, were designed using a web-based guide-RNA designer (http://www.rgenome.net/) and subsequently cloned into the Cas12i expression vector (a gift from Dr. Wei Li of the Institute of Zoology, Chinese Academy of Sciences) (*56*). To create CD33 knockout hPSCs (CD33^KO^-hPSCs), the two Cas12i-sgRNA expression vectors and a donor vector carrying the neomycin resistance gene (NeoR) were co-transfected into hPSCs using electroporation. Five days post-transfection, G418 (InvivoGen) was introduced to select the targeted hPSCs for one week. Following G418 selection, the cells were digested and single cells were seeded into 96-well plates to form single-cell clones. PCR and sequencing were used to verify the mutation sites, thereby establishing CD33 knockout clones.

### Genome PCR

Single-cell clones of hPSCs resistant to G418 were harvested for DNA extraction using the TIANamp Genomic DNA kit (TIANGEN). The CD33 locus target sequences were amplified with the 2 × Rapid Taq Master Mix (Vazyme) and the primer pairs depicted in **fig. S4**.

### AML xenograft models with CAR-iNK cell treatment

The MSLN^+^ HL-60 luci^+^ cells (5 × 10^5^ cells/mouse) were injected intravenously into NCG mice (NOD/ShiLtJGpt-*Prkdc*^em26Cd52^*Il2rg*^em26Cd22^/Gpt strain, GemPharmatech Co., Ltd.) to establish human AML xenograft models on day -1. On day 0, bioluminescence imaging (BLI, IVIS Spectrum PerkinElmer) was used to quantify the tumor graft burden in each group. After irradiation (1.0 Gy, Rad Source RS2000), the animals were randomly assigned to treatment groups and received an intravenous injection of the equivalent of 1 × 10^7^ fresh CAR^+^ iNK or unmodified iNK cells. Three days later (day 3), the animals received the second intravenous injection with the same cell count. Tumor aggressiveness was determined weekly by BLI. Mice with a significant tumor burden were humanely euthanized. All animal experiments were conducted under the Institutional Animal Care and Use Committee (IACUC) of the Institute of Zoology of the Chinese Academy of Sciences (IOZ-2021-182).

### Statistical analysis

All quantitative analysis were performed with SPSS (IBM SPSS Statistics 25). The Shapiro-Wilk normality test assessed the normal distribution of the data. Quantitative differences (mean ± SD) between samples were compared using two-tailed Student’s t-test, Mann-Whitney U test, one-way ANOVA test, and two-way ANOVA test. The *P values* were two-sided, and *P* < 0.05 was considered statistically significant. Survival curves for tumor-bearing models were plotted using the Kaplan-Meier method and compared to the treatment group using logarithmic rank (Mantel-Cox) tests. Statistical analysis was performed using the GraphPad Prism 9 software.

## Acknowledgments

We thank the staffs of SPF-grade animal facilities at the Institute of Zoology, Chinese Academy of Sciences for supplying animal care service.

## Author contributions

Y.W. and X.J.Z. designed and conducted all experiments, performed data analysis, and wrote the manuscript. Z.Q.W., Z.Y.X., F.Z., Y.H.L., and P.C.L. conducted some experiments and performed data analysis. Y.Q.L. and Q.T.W. performed scRNA-seq and data analysis. L.Q.Z., C.X.X., D.H.H., L.J.L., Y.P.Z., Q.Z., H.M.Q., Y.C., Y.Y.S., C.Y.Z., J.C.X., Y.Q.Z., J.X.W., T.J.W., M.Y.Z., W.Y.Y., and M.M.L. participated in multiple experiments. J.Y.W., X.F.Z., Q.C., X.D., W.B.Q., and A.B.L. discussed the data and manuscript. F.X.H. designed the project, designed and guided all experiments, scrutinized all experimental data, and wrote the manuscript. J.Y.W. revised the manuscript and provided the final approval of the manuscript.

## Data and materials availability

The scRNA-seq data of EGFP-NK, CD33 CAR-NK, MSLN CAR-NK, and Loop CAR-NK cells stimulated by Nomo-1 cells have been uploaded to the Genome Sequence Archive public database (HRA008163). Other relevant information or data is available from the corresponding authors upon reasonable request.

## Supplementary Materials

**Fig. S1.**
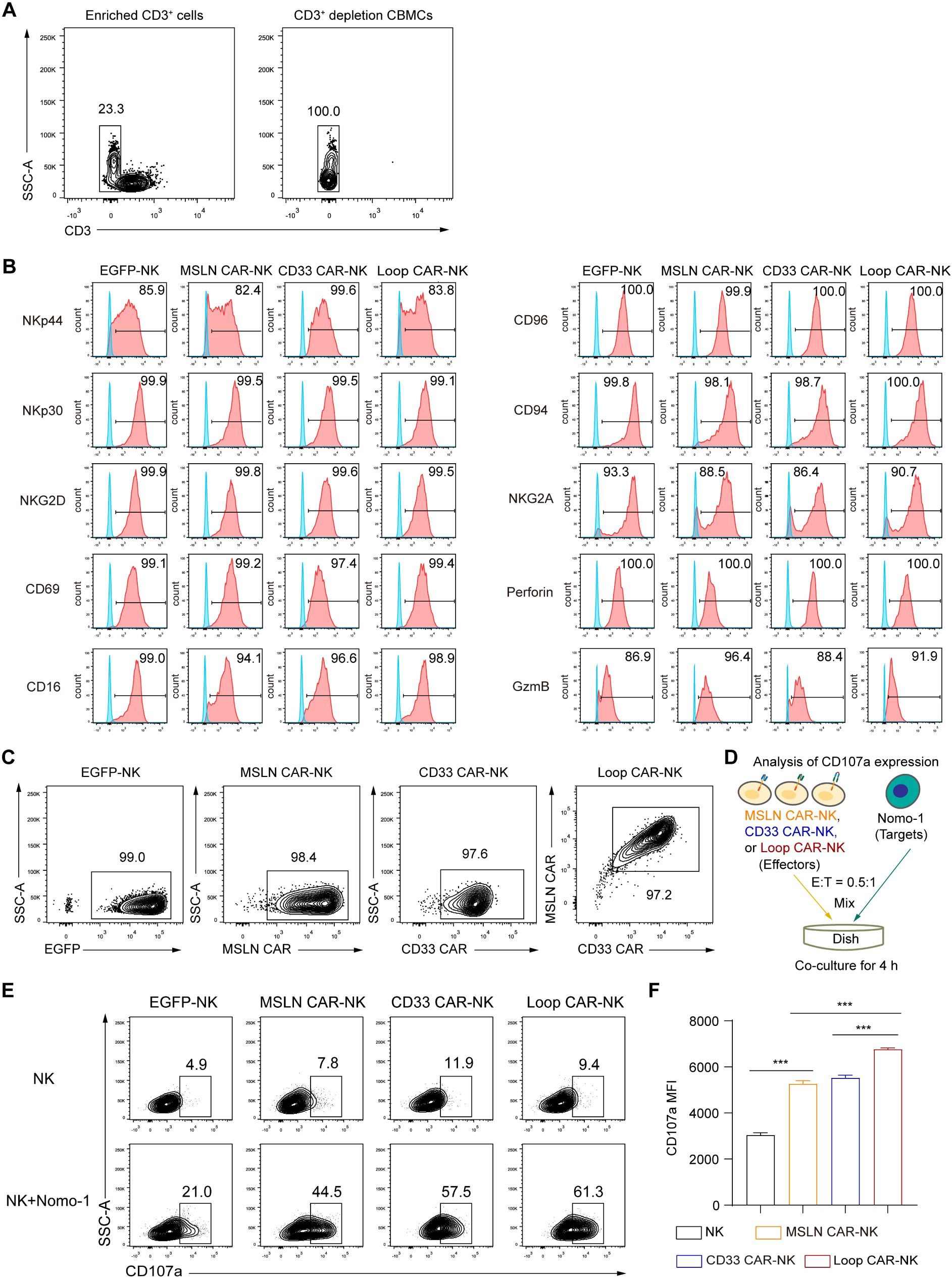
Phenotypes and effector function of Loop CAR-NK cells. (A) Flow cytometric analysis of the T cells (CD3^+^) residual in UCBMCs after CD3^+^ cells depletion on day -6. (B) Flow cytometry histograms showing the expression of typical NK cell receptors (NKp44, NKp30, NKG2D, CD69, CD16, CD96, CD94, and NKG2A) and cytotoxic granules (Perforin and GzmB) in respective NK cells. Control histograms represent FMO control. (C) The analysis of sorting purity for CD3^-^CD56^+^EGFP^+^ (EGFP-NK) and CD3^-^CD56^+^CAR^+^ (CAR-NK) cells post 24 h culturing. (D) Experimental design of the analysis of CD107a expression. EGFP-NK and CAR-NK cells were stimulated at an E:T ratio of 0.5:1 for 4 h. (E) Representative flow cytometry plots showing the expression of CD107a in EGFP-NK and CAR-NK cells in response to Nomo-1 cells. (F) Statistic analysis of the mean fluorescence intensity (MFI) of CD107a in (E), *n* = 3 repeats. Statistical analysis: one-way ANOVA, \*\*\**P* < 0.001.

**Fig. S2.**
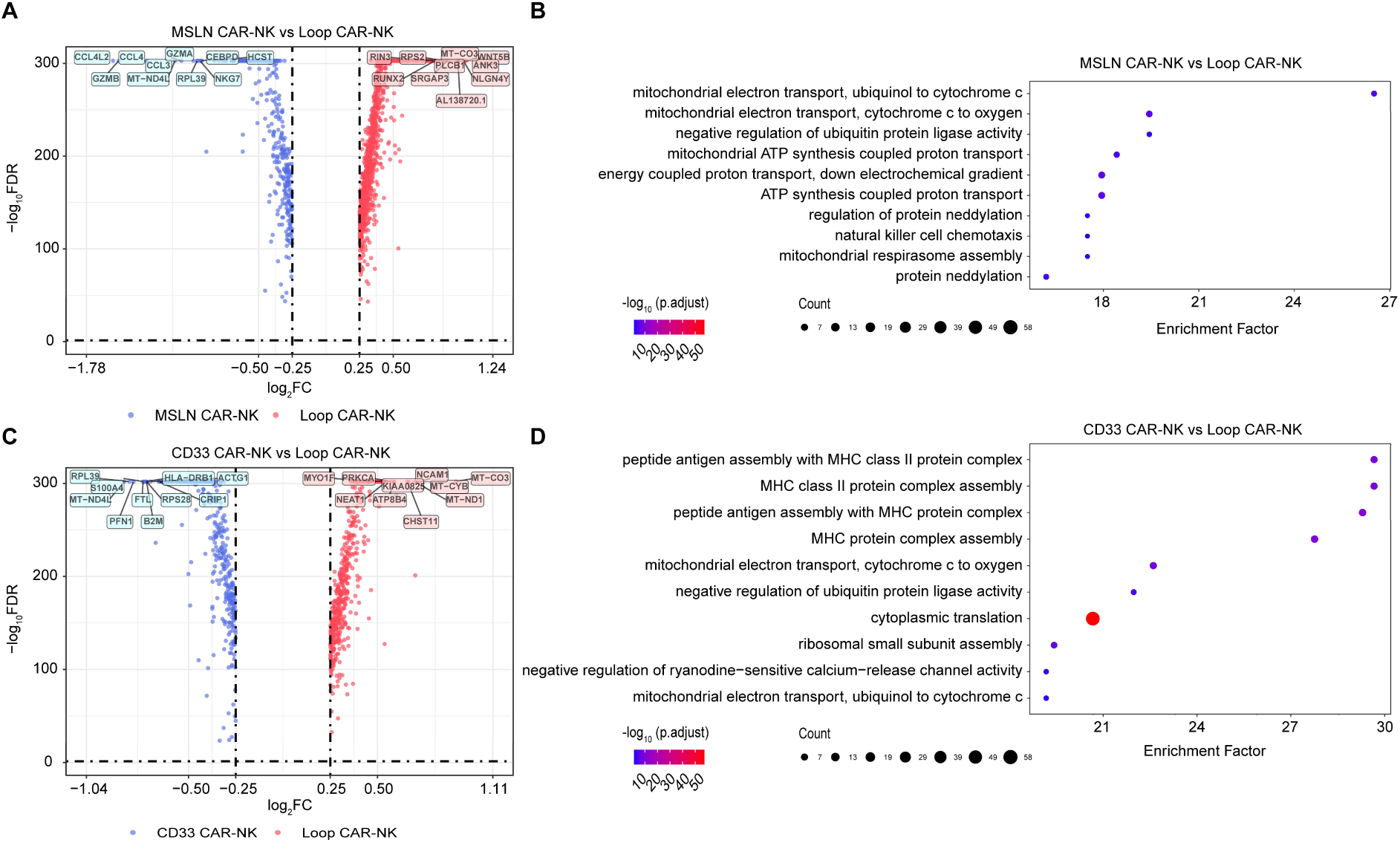
Differentially expressed genes of Loop CAR-NK cells compared with CD33 CAR-NK and MSLN CAR-NK cells. (A) Volcano plot of differentially expressed genes between Loop CAR-NK and MSLN CAR-NK cells. (B) GO analysis of differentially down-regulated genes in Loop CAR-NK cells compared with MSLN CAR-NK. Each data point represents a distinct GO set. Color scale corresponds to −log10 FDR (adjusted P values). Dot size was the number of genes enriched in one term. Enrichment Factor was the GeneRatio/BgRatio. (C) Volcano plot of differentially expressed genes between Loop CAR-NK and CD33 CAR-NK cells. (D) GO analysis of differentially down-regulated genes in Loop CAR-NK cells compared with CD33 CAR-NK cells. Each data point represents a distinct GO set. Color scale corresponds to −log10 FDR (adjusted P values). Dot size was the number of genes enriched in one term. Enrichment Factor was the GeneRatio/BgRatio.

**Fig. S3.**
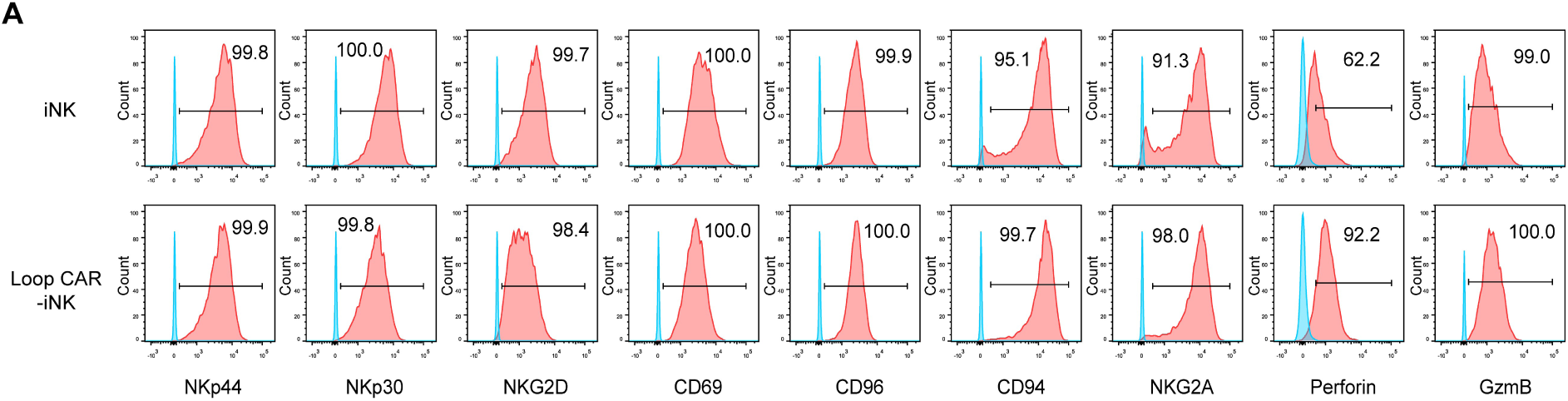
Analysis of typical NK molecules of Loop CAR-iNK cells. (A) Flow cytometry histograms showing the expression of typical NK cell receptors (NKp44, NKp30, NKG2D, CD69, CD96, CD94, and NKG2A) and cytotoxic granules (Perforin and GzmB) in respective iNK cells. Control histograms represent FMO control.

**Fig. S4.**
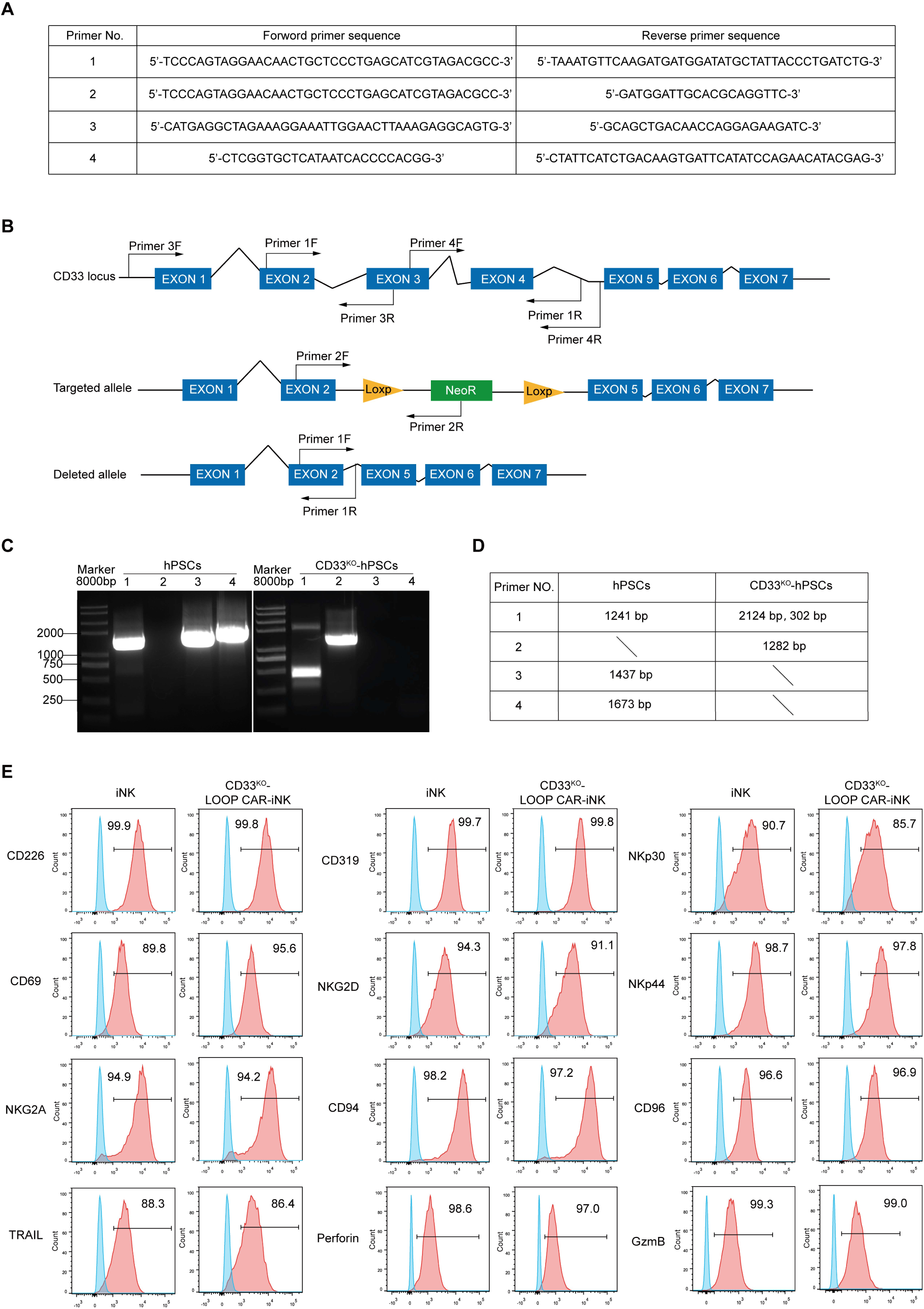
Identification of CD33 genotypes of CD33^KO^-hPSCs and molecular features of CD33^KO^-Loop CAR-iNK cells. (A) The sequences of four pairs of primers used for genotyping CD33 knock-out PSC clones. (B) Schematic diagram showing the location of primer pairs used for identifying CD33 genotypes. (C, D) PCR amplification results of each fragment using the indicated primer pairs. (E) Flow cytometry analysis of typical NK cell receptors (CD226, CD319, NKp30, CD69, NKG2D, NKp44, NKG2A, CD94, CD96, TARIL) and cytotoxic granules (Perforin and GzmB) in respective iNK cells. Control histograms represent FMO control.

**Table S1.**
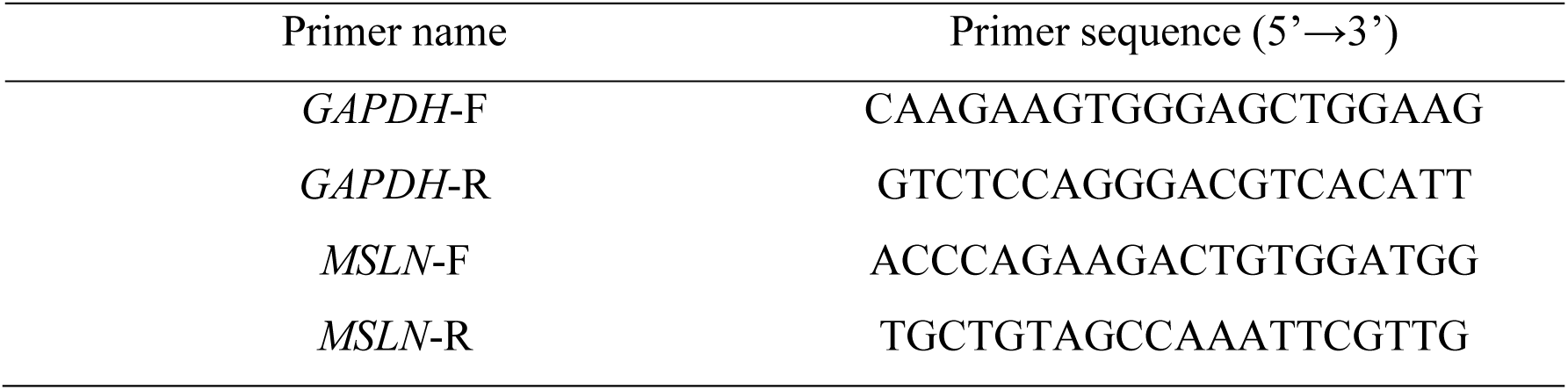
Primer sequences for real-time quantitative PCR of selected genes.

## References and Notes

1. H. Kantarjian, G. Borthakur, N. Daver, C. D. DiNardo, G. Issa, E. Jabbour, T. Kadia, K. Sasaki, N. J. Short, M. Yilmaz, F. Ravandi, Current status and research directions in acute myeloid leukemia. Blood Cancer J 14, 163 (2024).

2. D. Steinbach, A. Schramm, A. Eggert, M. Onda, K. Dawczynski, A. Rump, I. Pastan, S. Wittig, N. Pfaffendorf, A. Voigt, F. Zintl, B. Gruhn, Identification of a set of seven genes for the monitoring of minimal residual disease in pediatric acute myeloid leukemia. Clin Cancer Res 12, 2434–2441 (2006).

3. D. Steinbach, M. Onda, A. Voigt, K. Dawczynski, S. Wittig, R. Hassan, B. Gruhn, I. Pastan, Mesothelin, a possible target for immunotherapy, is expressed in primary AML cells. Eur J Haematol 79, 281–286 (2007).

4. A. J. Kaeding, S. P. Barwe, A. Gopalakrishnapillai, R. E. Ries, T. A. Alonzo, R. B. Gerbing, C. Correnti, M. R. Loken, L. E. Broderson, L. Pardo, Q. H. Le, T. Tang, A. R. Leonti, J. L. Smith, C. K. Chou, M. Xu, T. Triche, S. M. Kornblau, E. A. Kolb, K. Tarlock, S. Meshinchi, Mesothelin is a novel cell surface disease marker and potential therapeutic target in acute myeloid leukemia. Blood Adv 5, 2350–2361 (2021).

5. Q. Le, S. Castro, T. Tang, A. M. Loeb, T. Hylkema, C. N. McKay, L. Perkins, S. Srivastava, L. Call, J. Smith, A. Leonti, R. Ries, L. Pardo, M. R. Loken, C. Correnti, S. Fiorenza, C. J. Turtle, S. Riddell, K. Tarlock, S. Meshinchi, Therapeutic Targeting of Mesothelin with Chimeric Antigen Receptor T Cells in Acute Myeloid Leukemia. Clin Cancer Res 27, 5718–5730 (2021).

6. K. C. Hsu, C. A. Keever-Taylor, A. Wilton, C. Pinto, G. Heller, K. Arkun, R. J. O’Reilly, M. M. Horowitz, B. Dupont, Improved outcome in HLA-identical sibling hematopoietic stem-cell transplantation for acute myelogenous leukemia predicted by KIR and HLA genotypes. Blood 105, 4878–4884 (2005).

7. S. Z. D’Silva, M. Singh, A. S. Pinto, NK cell defects: implication in acute myeloid leukemia. Front Immunol 14, 1112059 (2023).

8. Y. Kaito, Y. Imai, Evolution of natural killer cell-targeted therapy for acute myeloid leukemia. Int J Hematol 120, 34–43 (2024).

9. Y. Masamoto, M. Kurokawa, A key to engineering natural killer cells to attack acute myeloid leukemia. Haematologica 109, 1032–1034 (2024).

10. R. Huang, X. Wang, H. Yan, X. Tan, Y. Ma, M. Wang, X. Han, J. Liu, L. Gao, L. Gao, G. Jing, C. Zhang, Q. Wen, X. Zhang, Safety and efficacy of CD33-targeted CAR-NK cell therapy for relapsed/refractory AML: preclinical evaluation and phase I trial. Exp Hematol Oncol 14, 1 (2025).

11. D. Huang, J. Li, F. Hu, C. Xia, Q. Weng, T. Wang, H. Peng, B. Wu, H. Wu, J. Xiong, Y. Lin, Y. Wang, Q. Zhang, X. Liu, L. Liu, X. Zheng, Y. Geng, X. Du, X. Zhu, L. Wang, J. Hao, J. Wang, Lateral plate mesoderm cell-based organoid system for NK cell regeneration from human pluripotent stem cells. Cell Discov 8, 121 (2022).

12. A. Eckstrom, A. Tyagi, S. Mahmood, L. Wong, B. Valamehr, A. Rao, A. Agrawal, M. Siddiqui, V. L. Battula, FT538, iPSC-derived NK cells, enhance AML cell killing when combined with chemotherapy. J Cell Mol Med 29, e70169 (2025).

13. D. A. Knorr, Z. Ni, D. Hermanson, M. K. Hexum, L. Bendzick, L. J. Cooper, D. A. Lee, D. S. Kaufman, Clinical-scale derivation of natural killer cells from human pluripotent stem cells for cancer therapy. Stem Cells Transl Med 2, 274–283 (2013).

14. Y. Li, D. L. Hermanson, B. S. Moriarity, D. S. Kaufman, Human iPSC-Derived Natural Killer Cells Engineered with Chimeric Antigen Receptors Enhance Anti-tumor Activity. Cell Stem Cell 23, 181–192 e185 (2018).

15. E. J. Orlando, X. Han, C. Tribouley, P. A. Wood, R. J. Leary, M. Riester, J. E. Levine, M. Qayed, S. A. Grupp, M. Boyer, B. De Moerloose, E. R. Nemecek, H. Bittencourt, H. Hiramatsu, J. Buechner, S. M. Davies, M. R. Verneris, K. Nguyen, J. L. Brogdon, H. Bitter, M. Morrissey, P. Pierog, S. Pantano, J. A. Engelman, W. Winckler, Genetic mechanisms of target antigen loss in CAR19 therapy of acute lymphoblastic leukemia. Nat Med 24, 1504–1506 (2018).

16. M. L. Cooper, J. Choi, K. Staser, J. K. Ritchey, J. M. Devenport, K. Eckardt, M. P. Rettig, B. Wang, L. G. Eissenberg, A. Ghobadi, L. N. Gehrs, J. L. Prior, S. Achilefu, C. A. Miller, C. C. Fronick, J. O’Neal, F. Gao, D. M. Weinstock, A. Gutierrez, R. S. Fulton, J. F. DiPersio, An “off-the-shelf” fratricide-resistant CAR-T for the treatment of T cell hematologic malignancies. Leukemia 32, 1970–1983 (2018).

17. M. Hejazi, C. Zhang, S. B. Bennstein, V. Balz, S. B. Reusing, M. Quadflieg, K. Hoerster, S. Heinrichs, H. Hanenberg, S. Oberbeck, M. Nitsche, S. Cramer, R. Pfeifer, P. Oberoi, H. Ruhl, J. Oldenburg, P. Brossart, P. A. Horn, F. Babor, W. S. Wels, J. C. Fischer, N. Moker, M. Uhrberg, CD33 Delineates Two Functionally Distinct NK Cell Populations Divergent in Cytokine Production and Antibody-Mediated Cellular Cytotoxicity. Front Immunol 12, 798087 (2021).

18. C. C. Kloss, M. Condomines, M. Cartellieri, M. Bachmann, M. Sadelain, Combinatorial antigen recognition with balanced signaling promotes selective tumor eradication by engineered T cells. Nat Biotechnol 31, 71–75 (2013).

19. D. Abate-Daga, M. L. Davila, CAR models: next-generation CAR modifications for enhanced T-cell function. Mol Ther Oncolytics 3, 16014 (2016).

20. D. Gomes-Silva, M. Srinivasan, S. Sharma, C. M. Lee, D. L. Wagner, T. H. Davis, R. H. Rouce, G. Bao, M. K. Brenner, M. Mamonkin, CD7-edited T cells expressing a CD7-specific CAR for the therapy of T-cell malignancies. Blood 130, 285–296 (2017).

21. M. Y. Kim, K. R. Yu, S. S. Kenderian, M. Ruella, S. Chen, T. H. Shin, A. A. Aljanahi, D. Schreeder, M. Klichinsky, O. Shestova, M. S. Kozlowski, K. D. Cummins, X. Shan, M. Shestov, A. Bagg, J. J. D. Morrissette, P. Sekhri, C. R. Lazzarotto, K. R. Calvo, D. B. Kuhns, R. E. Donahue, G. K. Behbehani, S. Q. Tsai, C. E. Dunbar, S. Gill, Genetic Inactivation of CD33 in Hematopoietic Stem Cells to Enable CAR T Cell Immunotherapy for Acute Myeloid Leukemia. Cell 173, 1439–1453 e1419 (2018).

22. N. Albinger, R. Pfeifer, M. Nitsche, S. Mertlitz, J. Campe, K. Stein, H. Kreyenberg, R. Schubert, M. Quadflieg, D. Schneider, M. W. M. Kuhn, O. Penack, C. Zhang, N. Moker, E. Ullrich, Primary CD33-targeting CAR-NK cells for the treatment of acute myeloid leukemia. Blood Cancer J 12, 61 (2022).

23. R. Handgretinger, H. J. Schafer, F. Baur, D. Frank, C. Ottenlinger, H. J. Buhring, D. Niethammer, Expression of an early myelopoietic antigen (CD33) on a subset of human umbilical cord blood-derived natural killer cells. Immunol Lett 37, 223–228 (1993).

24. M. Whitlow, B. A. Bell, S. L. Feng, D. Filpula, K. D. Hardman, S. L. Hubert, M. L. Rollence, J. F. Wood, M. E. Schott, D. E. Milenic, et al., An improved linker for single-chain Fv with reduced aggregation and enhanced proteolytic stability. Protein Eng 6, 989–995 (1993).

25. Y. Wang, J. Li, Z. Wang, Y. Liu, T. Wang, M. Zhang, C. Xia, F. Zhang, D. Huang, L. Zhang, Y. Zhao, L. Liu, Y. Zhu, H. Qi, X. Zhu, W. Qian, F. Hu, J. Wang, Comparison of seven CD19 CAR designs in engineering NK cells for enhancing anti-tumour activity. Cell Prolif 57, e13683 (2024).

26. S. O. Mathew, K. K. Rao, J. R. Kim, N. D. Bambard, P. A. Mathew, Functional role of human NK cell receptor 2B4 (CD244) isoforms. Eur J Immunol 39, 1632–1641 (2009).

27. A. Gutierrez-Guerrero, I. Mancilla-Herrera, J. L. Maravillas-Montero, I. Martinez-Duncker, A. Veillette, M. E. Cruz-Munoz, SLAMF7 selectively favors degranulation to promote cytotoxicity in human NK cells. Eur J Immunol 52, 62–74 (2022).

28. D. F. Barber, M. Faure, E. O. Long, LFA-1 contributes an early signal for NK cell cytotoxicity. J Immunol 173, 3653–3659 (2004).

29. Y. Wang, X. Tong, G. Li, J. Li, M. Deng, X. Ye, Ankrd17 positively regulates RIG-I-like receptor (RLR)-mediated immune signaling. Eur J Immunol 42, 1304–1315 (2012).

30. R. Narita, K. Takahasi, E. Murakami, E. Hirano, S. P. Yamamoto, M. Yoneyama, H. Kato, T. Fujita, A novel function of human Pumilio proteins in cytoplasmic sensing of viral infection. PLoS Pathog 10, e1004417 (2014).

31. J. T. Gunesch, A. L. Dixon, T. A. Ebrahim, M. M. Berrien-Elliott, S. Tatineni, T. Kumar, E. Hegewisch-Solloa, T. A. Fehniger, E. M. Mace, CD56 regulates human NK cell cytotoxicity through Pyk2. Elife 9, (2020).

32. D. Sancho, M. Nieto, M. Llano, J. L. Rodriguez-Fernandez, R. Tejedor, S. Avraham, C. Cabanas, M. Lopez-Botet, F. Sanchez-Madrid, The tyrosine kinase PYK-2/RAFTK regulates natural killer (NK) cell cytotoxic response, and is translocated and activated upon specific target cell recognition and killing. J Cell Biol 149, 1249–1262 (2000).

33. H. Ham, W. Huynh, R. A. Schoon, R. D. Vale, D. D. Billadeau, HkRP3 is a microtubule-binding protein regulating lytic granule clustering and NK cell killing. J Immunol 194, 3984–3996 (2015).

34. H. Ham, S. Guerrier, J. Kim, R. A. Schoon, E. L. Anderson, M. J. Hamann, Z. Lou, D. D. Billadeau, Dedicator of cytokinesis 8 interacts with talin and Wiskott-Aldrich syndrome protein to regulate NK cell cytotoxicity. J Immunol 190, 3661–3669 (2013).

35. Y. Liu, M. Zhang, X. Shen, C. Xia, F. Hu, D. Huang, Q. Weng, Q. Zhang, L. Liu, Y. Zhu, L. Wang, J. Hao, M. Zhang, T. Wang, J. Wang, Mesothelin CAR-engineered NK cells derived from human embryonic stem cells suppress the progression of human ovarian cancer in animals. Cell Prolif 57, e13727 (2024).

36. B. Xie, Z. Li, J. Zhou, W. Wang, Current Status and Perspectives of Dual-Targeting Chimeric Antigen Receptor T-Cell Therapy for the Treatment of Hematological Malignancies. Cancers (Basel) 14, (2022).

37. X. Wang, Z. Dong, D. Awuah, W. C. Chang, W. A. Cheng, V. Vyas, S. C. Cha, A. J. Anderson, T. Zhang, Z. Wang, S. J. Szymura, B. Z. Kuang, M. C. Clark, I. Aldoss, S. J. Forman, L. W. Kwak, H. Qin, CD19/BAFF-R dual-targeted CAR T cells for the treatment of mixed antigen-negative variants of acute lymphoblastic leukemia. Leukemia 36, 1015–1024 (2022).

38. Z. Chen, Y. Liu, N. Chen, H. Xing, Z. Tian, K. Tang, Q. Rao, Y. Xu, Y. Wang, M. Wang, J. Wang, Loop CD20/CD19 CAR-T cells eradicate B-cell malignancies efficiently. Sci China Life Sci 66, 754–770 (2023).

39. J. S. Orange, K. E. Harris, M. M. Andzelm, M. M. Valter, R. S. Geha, J. L. Strominger, The mature activating natural killer cell immunologic synapse is formed in distinct stages. Proc Natl Acad Sci U S A 100, 14151–14156 (2003).

40. E. Liu, Y. Tong, G. Dotti, H. Shaim, B. Savoldo, M. Mukherjee, J. Orange, X. Wan, X. Lu, A. Reynolds, M. Gagea, P. Banerjee, R. Cai, M. H. Bdaiwi, R. Basar, M. Muftuoglu, L. Li, D. Marin, W. Wierda, M. Keating, R. Champlin, E. Shpall, K. Rezvani, Cord blood NK cells engineered to express IL-15 and a CD19-targeted CAR show long-term persistence and potent antitumor activity. Leukemia 32, 520–531 (2018).

41. G. Alter, J. M. Malenfant, M. Altfeld, CD107a as a functional marker for the identification of natural killer cell activity. J Immunol Methods 294, 15–22 (2004).

42. M. Zhang, M. E. March, W. S. Lane, E. O. Long, A signaling network stimulated by beta2 integrin promotes the polarization of lytic granules in cytotoxic cells. Sci Signal 7, ra96 (2014).

43. T. Ulyanova, J. Blasioli, T. A. Woodford-Thomas, M. L. Thomas, The sialoadhesin CD33 is a myeloid-specific inhibitory receptor. Eur J Immunol 29, 3440–3449 (1999).

44. G. M. Robbins, M. Wang, E. J. Pomeroy, B. S. Moriarity, Nonviral genome engineering of natural killer cells. Stem Cell Res Ther 12, 350 (2021).

45. F. Cichocki, S. J. C. van der Stegen, J. S. Miller, Engineered and banked iPSCs for advanced NK- and T-cell immunotherapies. Blood 141, 846–855 (2023).

46. B. H. Goldenson, P. Hor, D. S. Kaufman, iPSC-Derived Natural Killer Cell Therapies - Expansion and Targeting. Front Immunol 13, 841107 (2022).

47. J. Liu, J. Tong, H. Yang, Targeting CD33 for acute myeloid leukemia therapy. BMC Cancer 22, 24 (2022).

48. N. N. Shah, J. Azzi, B. W. Cooper, A. Deol, J. DiPersio, D. Koura, B. McClune, L. S. Muffly, M. U. Mushtaq, R. Narayan, H. C. Suh, G. Yanik, S. Nath, J. Whangbo, G. Koehne, Phase 1/2 Study of Donor-Derived Anti-CD33 Chimeric Antigen Receptor Expressing T Cells (VCAR33) in Patients with Relapsed or Refractory Acute Myeloid Leukemia after Allogeneic Hematopoietic Cell Transplantation. Blood 142, 4862–4862 (2023).

49. C. Alewine, L. Xiang, T. Yamori, G. Niederfellner, K. Bosslet, I. Pastan, Efficacy of RG7787, a next-generation mesothelin-targeted immunotoxin, against triple-negative breast and gastric cancers. Mol Cancer Ther 13, 2653–2661 (2014).

50. P. Awuah, T. K. Bera, M. Folivi, O. Chertov, I. Pastan, Reduced Shedding of Surface Mesothelin Improves Efficacy of Mesothelin-Targeting Recombinant Immunotoxins. Mol Cancer Ther 15, 1648–1655 (2016).

51. P. R. Hamann, L. M. Hinman, I. Hollander, C. F. Beyer, D. Lindh, R. Holcomb, W. Hallett, H. R. Tsou, J. Upeslacis, D. Shochat, A. Mountain, D. A. Flowers, I. Bernstein, Gemtuzumab ozogamicin, a potent and selective anti-CD33 antibody-calicheamicin conjugate for treatment of acute myeloid leukemia. Bioconjug Chem 13, 47–58 (2002).

52. P. S. Chowdhury, J. L. Viner, R. Beers, I. Pastan, Isolation of a high-affinity stable single-chain Fv specific for mesothelin from DNA-immunized mice by phage display and construction of a recombinant immunotoxin with anti-tumor activity. Proc Natl Acad Sci U S A 95, 669–674 (1998).

53. T. J. Fry, N. N. Shah, R. J. Orentas, M. Stetler-Stevenson, C. M. Yuan, S. Ramakrishna, P. Wolters, S. Martin, C. Delbrook, B. Yates, H. Shalabi, T. J. Fountaine, J. F. Shern, R. G. Majzner, D. F. Stroncek, M. Sabatino, Y. Feng, D. S. Dimitrov, L. Zhang, S. Nguyen, H. Qin, B. Dropulic, D. W. Lee, C. L. Mackall, CD22-targeted CAR T cells induce remission in B-ALL that is naive or resistant to CD19-targeted CAR immunotherapy. Nat Med 24, 20–28 (2018).

54. I. Riviere, K. Brose, R. C. Mulligan, Effects of retroviral vector design on expression of human adenosine deaminase in murine bone marrow transplant recipients engrafted with genetically modified cells. Proc Natl Acad Sci U S A 92, 6733–6737 (1995).

55. K. B. Sanborn, G. D. Rak, A. N. Mentlik, P. P. Banerjee, J. S. Orange, Analysis of the NK cell immunological synapse. Methods Mol Biol 612, 127–148 (2010).

56. Y. Chen, Y. Hu, X. Wang, S. Luo, N. Yang, Y. Chen, Z. Li, Q. Zhou, W. Li, Synergistic engineering of CRISPR-Cas nucleases enables robust mammalian genome editing. Innovation (Camb) 3, 100264 (2022).

